# Sleep and sleep deprivation differentially alter white matter microstructure: a mixed model design utilising advanced diffusion modelling

**DOI:** 10.1101/2020.08.24.259432

**Authors:** Irene Voldsbekk, Inge Groote, Nathalia Zak, Daniël Roelfs, Oliver Geier, Paulina Due-Tønnessen, Lise-Linn Løkken, Marie Strømstad, Taran Y. Blakstvedt, Yvonne S. Kuiper, Torbjørn Elvsåshagen, Lars T. Westlye, Atle Bjørnerud, Ivan I. Maximov

**Affiliations:** Department of Psychology, University of Oslo, Oslo, Norway; Norwegian Centre for Mental Disorders Research (NORMENT), Oslo University Hospital, Oslo, Norway; Computational Radiology and Artificial Intelligence (CRAI), Division of Radiology and Nuclear Medicine, Oslo University Hospital, Oslo, Norway; Department of Diagnostic Physics, Division of Radiology and Nuclear Medicine, Oslo University Hospital, Oslo, Norway; Institute of Clinical Medicine, University of Oslo, Oslo, Norway; Division of Radiology and Nuclear Medicine, Oslo University Hospital, Oslo, Norway; Department of Neurology, Oslo University Hospital, Oslo, Norway; Department of Physics, University of Oslo, Oslo, Norway; Department of Health and Functioning, Western Norway University of Applied Sciences, Bergen, Norway

**Keywords:** White matter, DWI, MRI, Sleep, Sleep deprivation, Structural plasticity

## Abstract

Sleep deprivation influences several critical functions, yet how it affects human brain white matter (WM) is not well understood. The aim of the present work was to investigate the effect of 32 hours of sleep deprivation on WM microstructure compared to changes observed in a normal sleep-wake cycle (SWC). To this end, we utilised diffusion weighted imaging (DWI) including the diffusion tensor model, diffusion kurtosis imaging and the spherical mean technique, a novel biophysical diffusion model. 46 healthy adults (23 sleep deprived vs 23 with normal SWC) underwent DWI across 4 time points (morning, evening, next day morning and next day afternoon, after a total of 32 hours). Linear mixed models revealed significant group × time interaction effects, indicating that sleep deprivation and normal SWC differentially affect WM microstructure. Voxel-wise comparisons showed that these effects spanned large, bilateral WM regions. These findings provide important insight into how sleep deprivation affects the human brain.

## 1 Introduction

The significance of sleep is illustrated by the fact that sleep is evolutionary conserved across species (Siegel, 2009) and by the detrimental effects of sleep loss on health and cognition (Goel, Basner, Rao, & Dinges, 2013). Although the precise underlying mechanisms remain to be elucidated, sleep deprivation disrupts optimal information processing and learning, and is associated with several neurological and psychiatric diseases (Krause et al., 2017). Sleep is postulated to be regulated by two separable, yet interacting, processes: sleep homeostasis, by which increasing sleep pressure accumulates as a function of time spent awake, and the circadian rhythm, which is an intrinsic oscillating cycle of 24 hours regulated by exposure to daylight (Borbély, Daan, Wirz-Justice, & Deboer, 2016). However, it still remains unclear how sleep loss and the disruption of these sleep regulating processes affect the human brain.

Neuroimaging studies suggest that sleep deprivation affect many aspects of human brain structure and function. Sleep deprivation has been associated with increases in cerebral blood flow and reductions in metabolic rate and markers of neuronal synaptic activity in several brain regions (Elvsåshagen et al., 2019; M. L. Thomas et al., 2000, 2003). Moreover, studies find reductions in magnetic resonance imaging (MRI)-based measures of cortical thickness and functional connectivity after 24 hours of sleep deprivation (Elvsåshagen et al., 2017; Kaufmann et al., 2016), indicating that MRI is sensitive to the rapid effects of sleep deprivation on the human brain. However, only a few studies have focused on how sleep deprivation affects human brain white matter (WM). Diffusion tensor imaging (DTI)-based indices of diffusivity along axonal fibres exhibited significant reductions after sleep deprivation (Elvsåshagen et al., 2015), while another study found no effect of sleep deprivation on mean diffusivity across the whole WM (Bernardi et al., 2016). One study found increased WM volume after 24 hours of sleep deprivation and intense cognitive training (Bernardi et al., 2016), while another study found no volume change (Demiral et al., 2019). Moreover, recovery sleep the following day was found to reverse the effects of sleep deprivation on diffusivity changes in the cortex (Bernardi et al., 2016). These inconsistent results from previous studies might be due to differences in methodology used to assess brain changes, and differences in study design, including attempts to control factors that might affect sleep regulating processes. However, they collectively indicate that sleep deprivation might have a considerable impact on structure and function in widespread brain regions. Yet, how the effects of sleep deprivation compare with those of a normal sleep-wake-cycle (SWC) remains to be clarified.

Recent evidence also points to the influence of time-of-day (TOD) effects on various imaging-derived measures of human brain structure and function. These measures include total brain volume (Nakamura, Brown, Narayanan, Collins, & Arnold, 2015; Trefler et al., 2016) and apparent cortical thickness based on T_1_-weighted data (Elvsåshagen et al., 2017), regional cerebral blood flow based on arterial spin labelling (ASL) (Elvsåshagen et al., 2019; Hodkinson et al., 2014), and functional connectivity based on functional MRI (Hodkinson et al., 2014; Kaufmann et al., 2016). In diffusion-weighted imaging (DWI), we previously reported TOD effects in voxel-wise WM in two separate samples (Elvsåshagen et al., 2015; Voldsbekk et al., 2020), while others found TOD effects on voxel-wise whole brain measures (Jiang et al., 2014) and at the interface of grey matter (GM) and the cerebrospinal fluid (CSF) (C. Thomas et al., 2018). These findings suggest that TOD effects might represent important confounds in neuroimaging studies, and thus warrant further investigation.

While this has yet to be confirmed, TOD effects may be driven by the SWC and sleep regulating processes such as circadian and homeostatic rhythms. In the current study, we aimed to control for this by comparing the effect of 32 hours of sleep deprivation on the human brain with a normal SWC. To do this, we utilised a mixed design to investigate WM microstructure across 32 hours of either sleep deprivation or normal SWC in two groups of 23 healthy young adults using advanced DWI. Specifically, we employed the spherical mean technique (SMT) (Kaden, Kelm, Carson, Does, & Alexander, 2016), a novel biophysical model in which the diffusion signal is compartmentalised into differing pools of water. This model affords estimations of tissue microstructure in the intra-axonal and extra-axonal space, respectively, which provide more insight into tissue properties underlying diffusivity changes in WM, compared to conventional DTI. For comparative purposes, we also included estimations derived from the diffusion tensor model (Basser, Mattiello, & Le Bihan, 1994) and diffusion kurtosis imaging (DKI) (Jensen, Helpern, Ramani, Lu, & Kaczynski, 2005). Furthermore, we controlled for a number of *Zeitgeber* signals that might influence biological clocks, such as food intake, caffeine intake, physical activity and exposure to blue-emitting light. We hypothesised that sleep deprivation would lead to reductions in measures of WM diffusivity compared to the normal SWC group, while the normal SWC group would demonstrate TOD variations consistent with previous studies. Finally, we explored the associations between WM diffusivity changes and measures of sleep-wake characteristics such as chronotype, sleep habits and vigilant attention.

## 2 Material and Methods

### 2.1 Ethics statement

This study was approved by the Regional Committee for Medical and Health Research Ethics, South-Eastern Norway (REK Sør-Øst, ref: 2017/2200) and conducted in line with the Declaration of Helsinki of 2013. Prior to participation, all participants gave their written informed consent. Participants received NOK 1000 for their participation.

### 2.2 Participants

The recruitment procedure and sample are described in earlier work (Voldsbekk et al., 2020). Healthy volunteers were recruited through adverts in a national newspaper and on social media. The recruitment process is illustrated in Figure 1. 127 volunteers were clinically screened by phone interview in which exclusion criteria were: history of any psychiatric disorder, severe or chronic somatic disorder, current intake of any regular medication, smoking, contraindications to MRI or living more than one hour of travel away from the MRI facility. This procedure excluded 41 volunteers. Of the remaining 86 volunteers, 15 withdrew from the study and 22 were cancelled due to logistic reasons. In addition, one participant was excluded due to illness during the study and two due to claustrophobic reaction in the scanner. The final sample consisted of 46 healthy adults (age 25.9 ± 6.9 years; 29 women).

**Figure 1.**
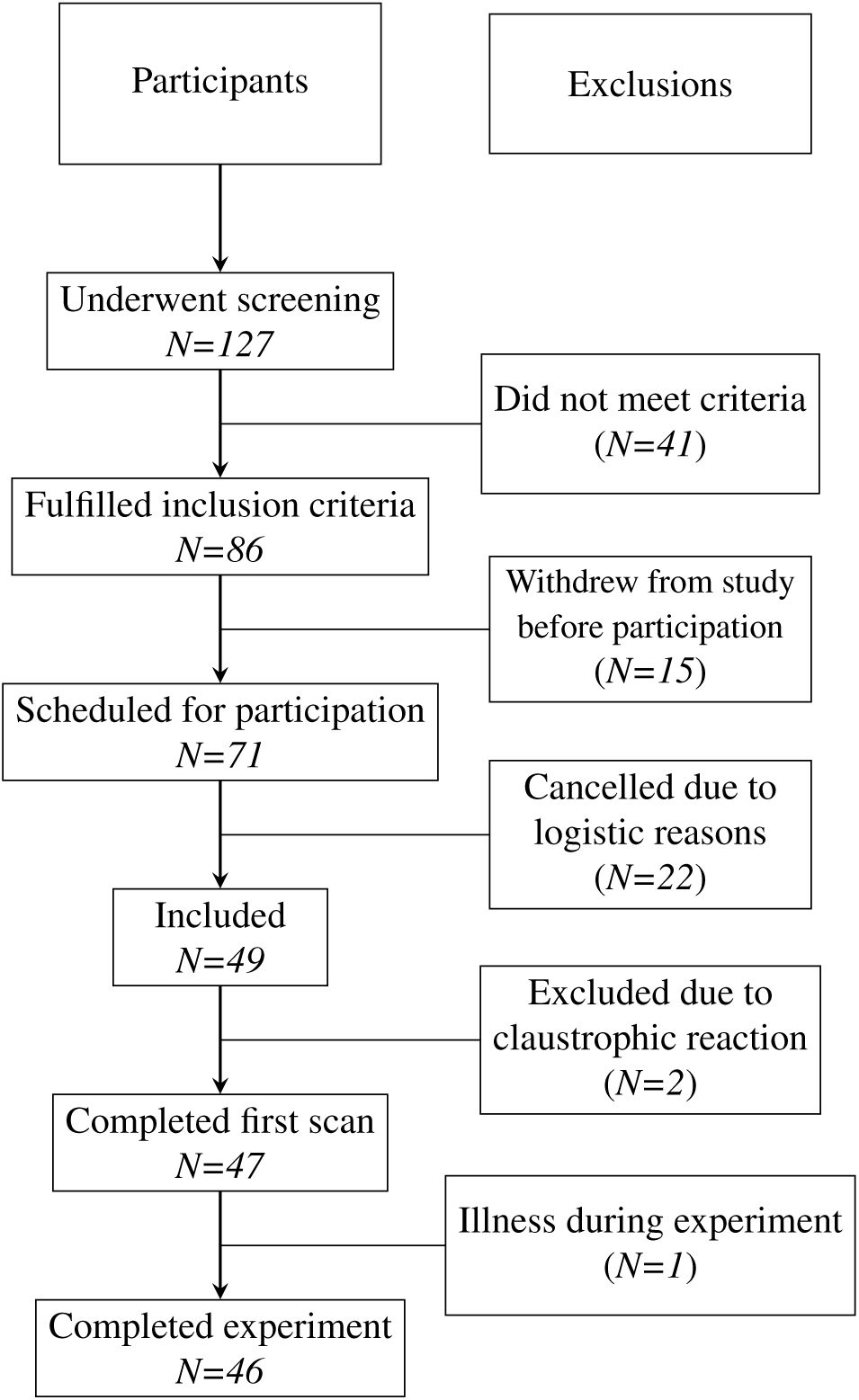
Overview of the inclusion process. 127 were screened and 86 of these met the inclusion criteria. 71 were then scheduled for participation but only 49 participated due to logistic reasons. Of these, two were excluded due to claustrophobic reaction and one due to illness during the experiment, which resulted in a final sample of 46.

### 2.3 Study design

Figure 2 presents an overview of the study design. Participants were scanned the first time fasting in the morning upon arrival at Oslo University Hospital, Rikshospitalet after a night of normal sleep in their own home (around 9 AM; TP1). The second scan took place in the evening approximately 11 hours later (around 8 PM; TP2), the third scan the next morning after an additional 12 hours (around 8 AM; TP3) and the final scan the same afternoon (around 4 PM; TP4). After the evening scan (TP2), participants were randomised by draw to either the sleep or the normal SWC condition. This was to ensure no difference in treatment from the experimenters or own behaviour during the first study day. Participants in the normal SWC (*n*=23) then went home to sleep, while those in the sleep deprivation group (*n*=23) stayed at the hospital. During their time at the hospital, participants followed a standardised activity protocol under continuous supervision by a research assistant to ensure that none fell asleep (see Voldsbekk et al., 2020 for a detailed description). In order to ensure that no one fell asleep during the MRI scans, a camera inside the scanner bore was used to monitor the eyes of the participants (Model 12M-i, MRC Systems GmbH, Heidelberg, Germany). The camera had a sampling rate of 25 frames per second and a resolution of 853×480 pixels. After each scan and every other hour, participants indicated their subjective sleepiness and every third hour they completed a measure of their alertness (see below). In addition, participants completed a battery of questionnaires regarding their health and sleep habits. Data was collected during the summer season of 2018. During this period, the daytime duration in Oslo, Norway was 17:46:29 ± 1:28:24 hours per day (Time and Date, 2020).

**Figure 2.**
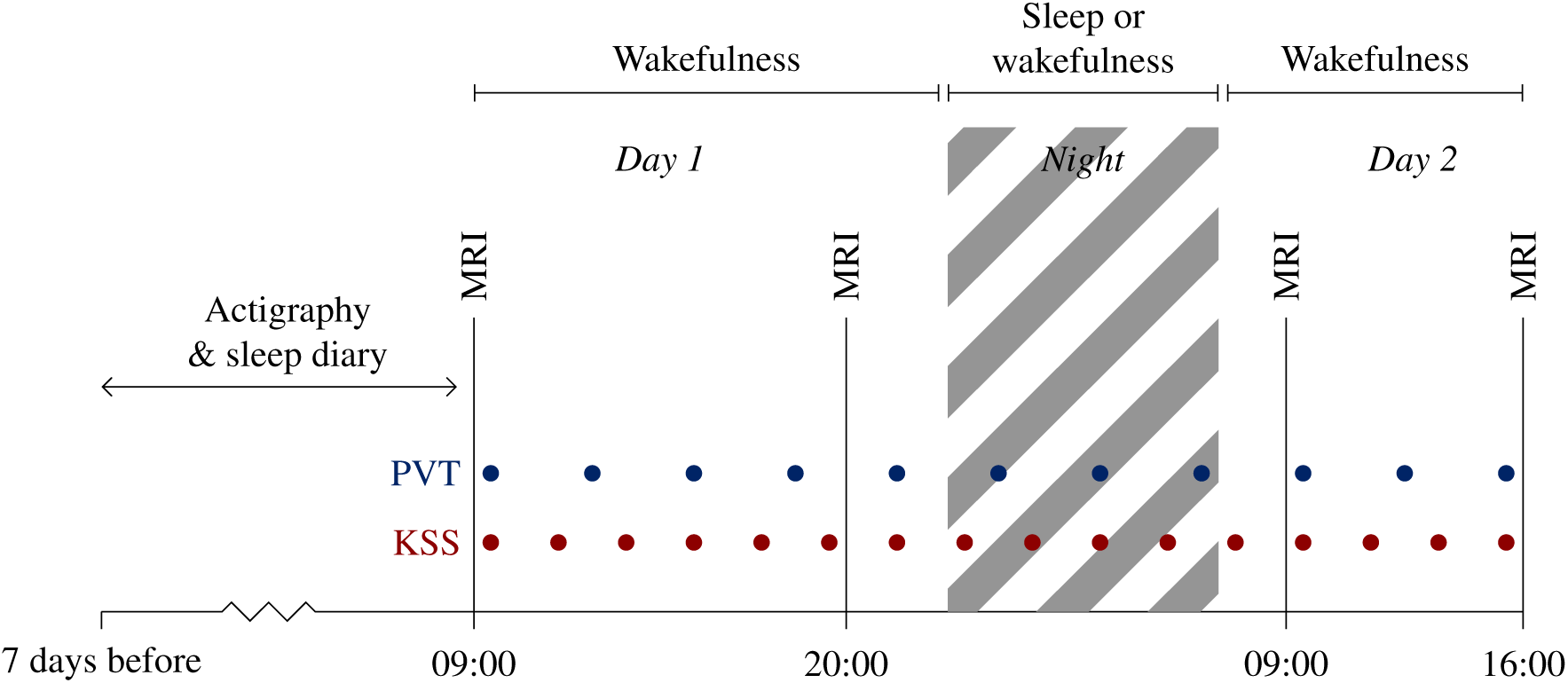
Overview of the study protocol. Participants underwent MRI in the morning after a night of sleep (time point (TP)1), in the evening after a day of waking (TP2), after a night of either sleep or sleep deprivation (TP3) and then in the afternoon on the second day (TP4). During the study, participants underwent tests of sleepiness (KSS) and alertness (PVT) every second and third hour, respectively. Seven days prior to the first MRI scan, participants underwent measurements of sleep-habits by actigraphy and self-report sleep diary.

### 2.4 Assessment of sleep habits

Sleep pattern prior to participation was recorded to ensure that participants had slept between six and nine hours each night in the preceding week. Participants in the normal SWC group also recorded their sleep pattern during the night of study in which they went home to sleep. Self-report data was obtained using a sleep diary (Bjorvatn, 2018) consisting of a 10-item semi-structured scale to be completed on a daily basis. It characterises sleep-related behaviour and quality, such as bedtime, rise time, perceived amount of time taken to fall asleep, sleep quality, duration of nightly awakenings if any, consumption of any sleep medication or other sleeping aid. We modified the scale to also include two items on caffeine intake and nicotine intake. In addition, we obtained behavioural data using a Condor Instruments ActTrust actigraph (São Paulo, Brazil). The actigraph measures an individual’s movements by a digital tri-axial accelerometer with a 60s epoch. The acquired data is then used to estimate sleep and circadian parameters. We combined the data from the sleep diaries and actigraphs in order to estimate the sleep patterns of each participant.

Moreover, participants were instructed to complete five standardised questionnaires regarding their sleep habits: the Bergen Insomnia Scale (Pallesen, Bjorvatn, Nordhus, Sivertsen, & Hjørnevik, 2008), the Epworth Sleepiness Scale (ESS) (Johns, 1991), the Global Sleep Assessment Questionnaire (GSAQ) (Roth et al., 2002), the Pittsburgh Sleep Quality Index (PSQI) (Buysse et al., 1989) and the Horne-Østberg Morningness Eveningness Questionnaire (MEQ) (Horne & Östberg, 1976). These questionnaires measure insomnia-related symptoms, daytime sleepiness, prevalence of sleep complaints, sleep quality, and chronotype, respectively.

### 2.5 Assessment of acute sleepiness and alertness

During wakefulness of the two groups, we obtained measures of subjective acute sleepiness and alertness every third hour. Participants filled out the Karolinska Sleepiness Scale (KSS), which is a 1-item self-report indication of sleepiness on a nine-point Likert scale (åkerstedt & Gillberg, 1990). To measure alertness, participants completed a computerised psychomotor vigilance test (PC-PVT) (Khitrov et al., 2014), in which the reaction time to a visual stimulus is recorded as a measurement of sustained attention (Dinges & Powell, 1985). Specifically, participants click a mouse button in response to a five-digit millisecond counter presented as the visual stimulus on a screen with random intervals. Performance on the task is quantified based on reaction time and lapses in the response, which is strongly correlated with sleep need and sleep deprivation (Basner & Dinges, 2011). For the current study, minor lapses were used as a measure of alertness. The PC-PVT was run for 10 minutes with Matlab 2017a (MathWorks, Massachusetts, USA) on a Lenovo laptop V510-15IKB with Windows 10 Pro using a Cooler Master gaming mouse model SGM-1006-KSOA1. The laptop display had a refresh rate of 60 Hz. The movement resolution of the mouse was 2000 dpi.

### 2.6 acquisition

Imaging was performed on a 3.0 Tesla Siemens Magnetom Prisma scanner (Siemens Healthcare, Erlangen, Germany) using a 32-channel head coil (Erlangen, Germany). The scan protocol consisted of a monoplanar diffusion encoding sequence with a single-shot echo-planar imaging (EPI) readout module (Setsompop et al., 2012). 76 axial slices with *b*-values=[500-1000-2000-3000](s/mm^2^) and non-coplanar diffusion-sensitised gradient directions (Stirnberg, Stöcker, & Shah, 2009) with the corresponding numbers of gradient directions n_dir_=[12-30-40-50] were acquired. The following parameters were applied: repetition time /echo time = 2450ms/85ms, field-of-view = 212 x 212mm^2^, slice thickness = 2mm, matrix = 106, voxel size = 2 × 2 × 2 mm, flip-angle = 78°, multi-band acceleration factor = 4. Acquisition time was 8 min 21 s. Five images of opposite phase-encode direction and b = 0 were also obtained for correction of susceptibility distortions with an acquisition time of 31 s.

### 2.7 Assessment of hydration and head motion

Immediately after each scan, a blood sample was drawn from the cubital vein for analysis of haematocrit, as a measure of hydration level. Haematocrit at each TP was included as a confounder in MRI analyses. In addition, an index of head motion was estimated based on displacement and rotation matrices obtained from the MRI scan. From these the framewise displacement (FD) was calculated (Power, Barnes, Snyder, Schlaggar, & Petersen, 2012), which was also included as a confounder in MRI analyses.

### 2.8 MRI preprocessing

DWI data were pre-processed and corrected using an optimised pipeline consisting of six steps (Maximov, Alnaes, & Westlye, 2019), including Rician noise correction (Veraart, Fieremans, & Novikov, 2016; Veraart, Novikov, et al., 2016), Gibbs ringing artefact correction (Kellner, Dhital, Kiselev, & Reisert, 2016), susceptibility distortion correction (Andersson, Skare, & Ashburner, 2003), eddy current correction (Andersson et al., 2017; Andersson, Graham, Zsoldos, & Sotiropoulos, 2016; Andersson & Sotiropoulos, 2016), field non-uniformity correction (Tustison et al., 2010) and smoothing using a 1mm^3^ Gaussian kernel. Metrics derived from DTI (Basser et al., 1994) and DKI (Jensen et al., 2005) were estimated using Matlab R2014a (MathWorks, Natick, Massachusetts, USA) as proposed by Veraart and colleagues (Veraart, Sijbers, Sunaert, Leemans, & Jeurissen, 2013). DTI metrics include the mean diffusivity (MD), a measure of mean diffusivity within a voxel; fractional anisotropy (FA), a measure of the directionality or anisotropy of the diffusion in each voxel; axial diffusivity (AD), a measure of diffusion parallel to the principal diffusion direction, and radial diffusivity (RD), a measure of diffusion perpendicular to the principal diffusion direction. In addition, DKI metrics include the mean kurtosis (MK), axial kurtosis (AK) and radial kurtosis (RK), which are estimations of the degree of deviation from a Gaussian diffusion probability distribution (Jensen et al., 2005). From the SMT model (Kaden et al., 2016) we estimated the intra-neurite volume fraction (INVF), a volume fraction of intra-axonal space relative to extra-axonal space; extra-neurite mean diffusivity (exMD), which is mean diffusivity in extra-axonal space; and extra-neurite radial diffusivity (exRD), which is a measure of diffusivity in extra-axonal space in the radial or transverse direction. Diffusion scalar maps were processed and prepared for hypothesis testing using tract-based spatial statistics (TBSS) (Smith et al., 2006). In this procedure, each map is aligned to standard space using non-linear registration and then thinned to create a mean skeleton based on fractional anisotropy (FA) metrics. A mean FA skeleton image across all participants was created, thresholded and binarised at 0.25 to reduce partial volume effects. Aligned image sets from each participant were then projected onto this mean skeleton before voxel-wise group-level comparisons. Mean values of diffusion metrics across the whole skeleton were calculated and further analysed in R (R Core Team, 2018).

### 2.9 Statistical analysis of diffusion data

To test for changes in diffusion data across 32 hours of sleep deprivation vs. normal SWC, linear mixed models were tested using the *lme* function in R (Bates & Pinheiro, 1998). For each diffusion metric, we tested for interactions between group (sleep deprivation or normal SWC) and time (TP1, TP2, TP3, TP4). In fitting the model, we scaled each variable and entered time and group as fixed effects. Participant ID was entered as a random effect. The model equation is shown in Eq. (1).

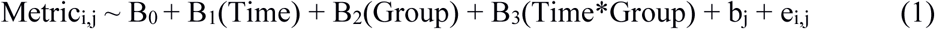

where

- Metric_i,j_ is the mean diffusion metric value for participant *i* at TP *j*;
- Time is the TPs;
- Group is the normal SWC or sleep deprivation);
- Time*Group is the interaction between the group term and the TP;
- B_0,1,2,3_ are the fixed effect coefficients: B_0_ is the intercept; B_1_ is the linear slope for TP; B_2_ is the linear slope for Group; B_3_ is the coefficient for the interaction term (Time*Group);
- b_j_ is the random effect coefficient for participant *i*; and
- e_i_, *j* is the error for participant *i* at TP *j*.

Visual inspection of residual plots did not reveal any large deviations from homoscedasticity or normality. Estimated marginal means (EMMs) were obtained with predictions from the fitted model using the *effect* function in R (Fox, 2003). To investigate any significant interactions in more detail, we then ran corresponding pairwise comparisons using the *emmeans* function in R (Lenth, Singmann, Love, Buerkner, & Herve, 2018). We ran 12 paired t-tests comparing mean diffusion values between two differing TPs separately for the sleep deprived and normal SWC group. In addition, we ran four independent t-tests comparing mean diffusion value between the sleep deprived and the normal SWC group at each TP. The significance threshold was set at *p* < 0.05, and the results were corrected for multiple comparisons using the false discovery rate (FDR) (Benjamini & Hochberg, 1995). To control for potential confounding factors, we reran the linear mixed model including age, sex, amount of head motion and haematocrit as covariates.

Next, group × time interaction effects in diffusion data were further investigated by voxel-wise tests using FSL tool randomise (Winkler, Ridgway, Webster, Smith, & Nichols, 2014). For each participant, the difference between two TPs was calculated by subtracting one skeleton map from the other. These difference images were then compared across groups by two sample t-tests and permutation testing. These tests were computed using threshold-free cluster enhancement (TFCE; Smith & Nichols, 2009) and 5000 permutations. Inherent to this method is a correction for family-wise error (FWE) across space.

### 2.10 Statistical analysis of sleep-wake characteristics

Pearson correlation analyses were conducted to test for associations between changes in diffusion metrics and sleep-wake-characteristics. In these analyses, the following measures were included as sleep-wake-characteristics: 1) change in sleepiness, as measured by KSS; 2) change in alertness, as measured by minor lapses on the PC-PVT; 3) chronotype, as measured by MEQ; 4) mean total sleep time (TST) per night for the previous week and 5) TST for the previous night, as measured by sleep diary and actigraphy. In addition, we conducted two additional linear mixed models to investigate group × time interaction effects on sleepiness and alertness, as measured by KSS and PC-PVT throughout the study.

## 3 Results

### 3.1 Sleep pattern assessment

All participants had slept approximately seven hours each night for the past week, as recorded by self-report and corroborated by actigraphy measurements (see Table 1). For the normal SWC group, there was no significant difference in total sleep time on the night between the first and second day of the study, as compared to the night prior to study start (t_(27)_ = -.31, *p* = .76). Across groups, participants slept significantly shorter on the night before the study as compared to their weekly average (t_(79)_ = 2.28, *p* = .03). A summary of sleep habits is reported in Table 1.

**Table 1.** Descriptive statistics for sleep characteristics.

| | All participants<br>($n = 46$) | Normal SWC<br>($n = 23$) | Sleep deprived<br>($n = 23$) |
| --- | --- | --- | --- |
| Age | 25.65 ± 6.73 | 25.8 ± 6.9 | 25.5 ± 6.7 |
| Sex (no. of women) | 29 (63%) | 15 (65%) | 14 (61%) |
| PSQI (0-42)<sup>a</sup> | 4.42 ± 1.9 | 4.55 ± 1.9 | 4.3 ± 1.94 |
| MEQ (16-86)<sup>a</sup> | 49.54 ± 9.07 | 49.6 ± 10.2 | 49.5 ± 8.02 |
| BIS (0-42)<sup>a</sup> | 7.12 ± 5.18 | 8.02 ± 6.22 | 6.26 ± 3.88 |
| ES (0-24)<sup>a</sup> | 6.44 ± 3.74 | 6.55 ± 4.52 | 6.35 ± 2.9 |
| GSAQ (15-60)<sup>a</sup> | 19.53 ± 2.48 | 20.2 ± 2.79 | 18.9 ± 1.98 |
| Average TST week before study<sup>1,b</sup> (h:min) | 6:58 ± 0:57 | 7:05 ± 0:55 | 6:51 ± 0:59 |
| Average TST night before study<sup>1,c</sup> (h:min) | 6:28 ± 1:02 | 6:28 ± 0:45 | 6:28 ± 1:16 |
| Average TST the night during the study | - | 6:32 ± 0:47 | - |

*Note.* PSQI; Pittsburgh Global Sleep Quality Index. MEQ; Horne-Östberg Morningness Eveningness Questionnaire. BIS; Bergen Insomnia Scale. ES; Epworth Sleepiness Scale. GSAQ; Global Sleep Assessment Questionnaire. TST; total sleep time. SWC; sleep-wake cycle.
<sup>1</sup> Reported total sleep time as measured by actigraphy and controlled against sleep diary.
<sup>a</sup> Missing for one participant.
<sup>b</sup> Missing for two participants.
<sup>c</sup> Missing for six participants.

### 3.2 Effect of sleep deprivation and normal sleep-wake on white matter microstructure

Results from the linear mixed model revealed significant main effects of time in INVF and exRD, but not exMD (see Table 2). Further, there were significant group × time interaction effects in exMD at TP3 and TP4, but not INVF and exRD. Figure 3 shows the EMMs with corresponding standard errors (SE) across time for each group. The group × time interaction effects in exMD at TP3 and TP4 remained significant after controlling for confounding factors age, sex, head motion and haematocrit (β = -.33, SE = .13, *p* = .03 and β = -.37, SE = .13, *p* = .02; FDR-corrected, respectively), indicating that the covariates could not fully explain this relationship. The main effects of time in INVF and exRD did not remain significant (*p* = .06 and *p* = .08; FDR-corrected, respectively). In DTI metrics, the linear mixed model revealed a main effect of time in FA, MD and RD, but no significant group × time interaction effects (See Supplementary Material 1 for tables and figures). In DKI metrics, there were a main effect of time in AK, but otherwise no significant effects (See Supplementary Material 2 for tables and figures).

**Table 2.**
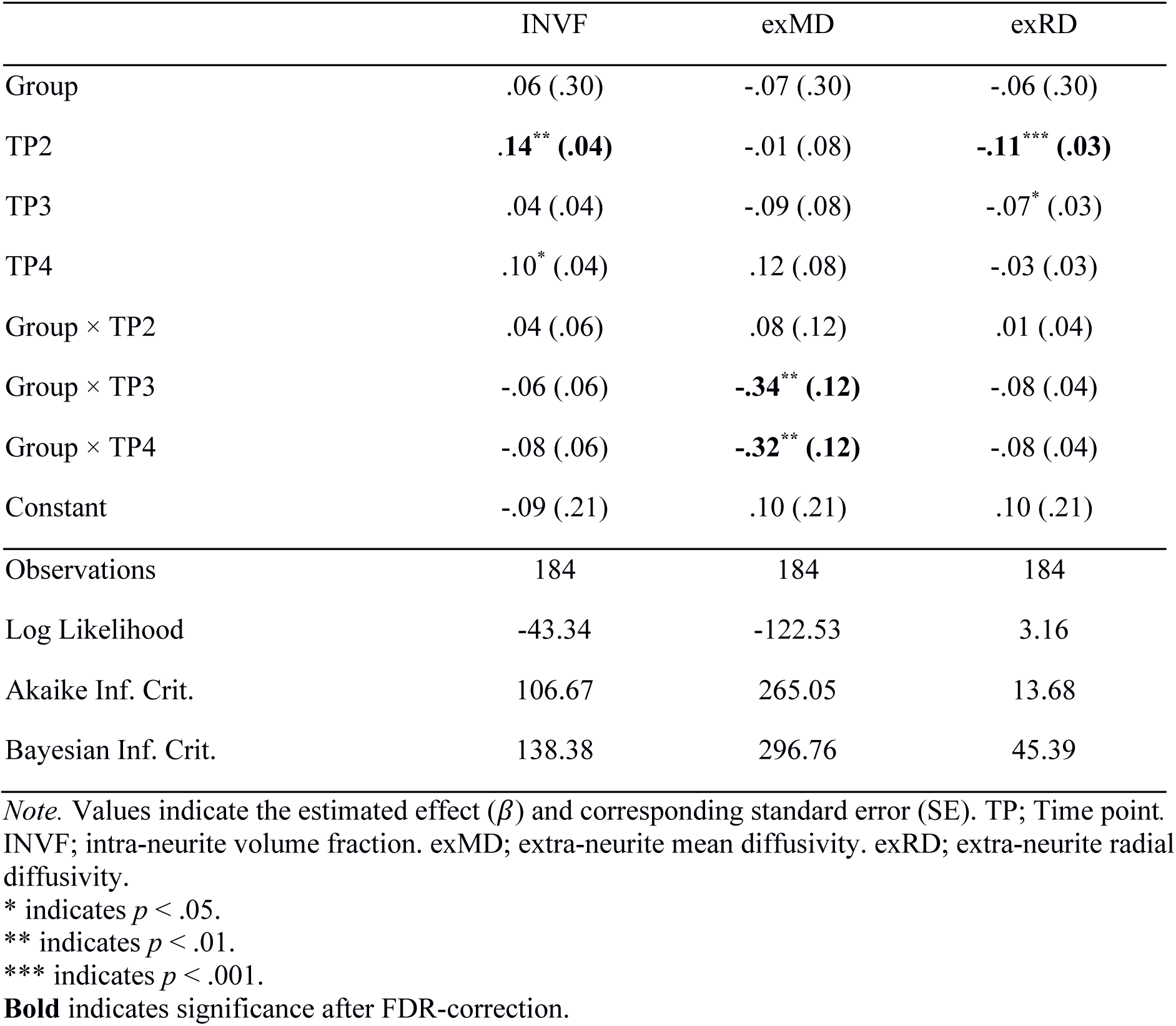
Linear mixed models for each SMT metric.

|  | INVf | exMD | exRD |
| --- | --- | --- | --- |
| Group | .06 (.30) | -.07 (.30) | -.06 (.30) |
| TP2 | **.14\*\* (.04)** | -.01 (.08) | **-.11\*\*\* (.03)** |
| TP3 | .04 (.04) | -.09 (.08) | -.07* (.03) |
| TP4 | .10* (.04) | .12 (.08) | -.03 (.03) |
| Group × TP2 | .04 (.06) | .08 (.12) | .01 (.04) |
| Group × TP3 | -.06 (.06) | **-.34\*\* (.12)** | -.08 (.04) |
| Group × TP4 | -.08 (.06) | **-.32\*\* (.12)** | -.08 (.04) |
| Constant | -.09 (.21) | .10 (.21) | .10 (.21) |
| Observations | 184 | 184 | 184 |
| Log Likelihood | -43.34 | -122.53 | 3.16 |
| Akaike Inf. Crit. | 106.67 | 265.05 | 13.68 |
| Bayesian Inf. Crit. | 138.38 | 296.76 | 45.39 |

*Note.* Values indicate the estimated effect ( $\beta$ ) and corresponding standard error (SE). TP; Time point. INVf; intra-neurite volume fraction. exMD; extra-neurite mean diffusivity. exRD; extra-neurite radial diffusivity.
\* indicates $p < .05$ .
\*\* indicates $p < .01$ .
\*\*\* indicates $p < .001$ .
**Bold** indicates significance after FDR-correction.

**Figure 3.**
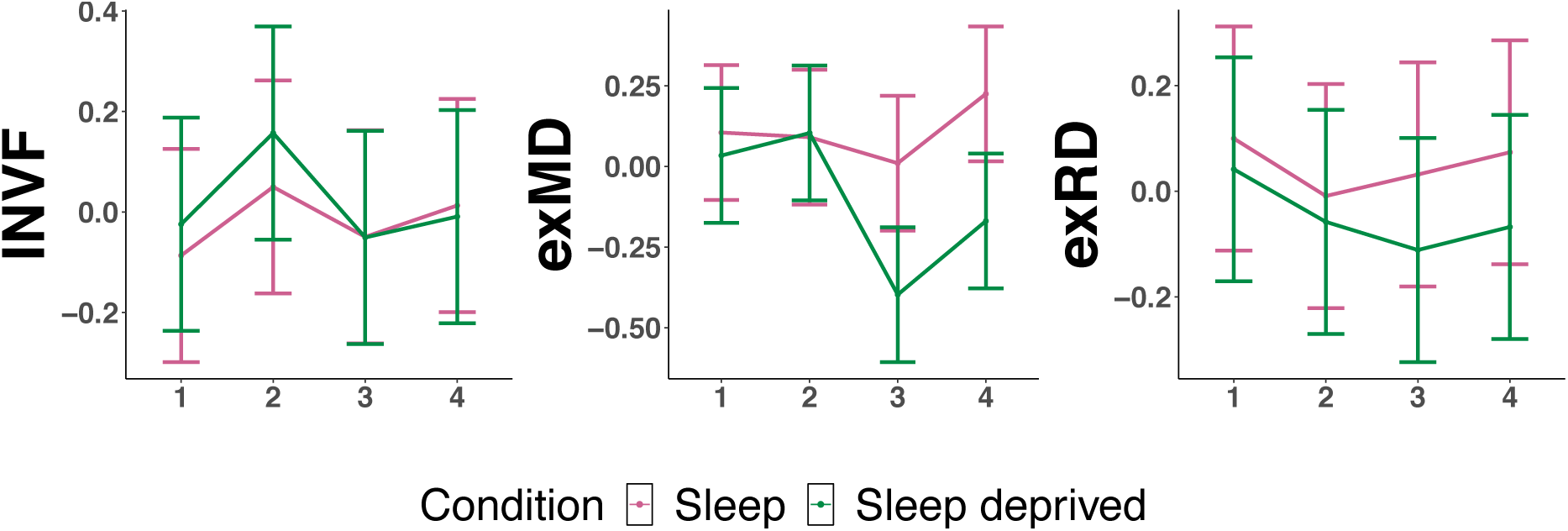
Estimated marginal means (y-axis) for the normal sleep-wake cycle group (pink) and sleep deprived group (green) across time points (x-axis). INVF; intra-neurite volume fraction. exMD; extra-neurite mean diffusivity. exRD; extra-neurite radial diffusivity.

#### 3.2.1 Pairwise comparisons of simple effects

The pairwise comparisons of simple effects in exMD showed significant decreases in the sleep deprived group from TP1 to TP3 (β = .43, SE = .08, *p* < .001; FDR-corrected), from TP1 to TP4 (β = .20, SE = .04, *p* = .01; FDR-corrected), from TP2 to TP3 (β = .50, SE = .08, *p* < .001; FDR-corrected) and from TP2 to TP4 (β = .27, SE = .04, *p* = .001; FDR-corrected). In addition, the sleep deprived group showed significant increases in exMD from TP3 to TP4 (β = .22, SE = .04, *p* < .01; FDR-corrected). The sleep group showed no significant pairwise comparisons in exMD after FDR-correction.

#### 3.2.2 Voxel-wise group × time interaction effects

Voxel-wise tests of group × time interaction effects revealed clusters with significant interactions in INVF, exMD and exRD from TP2 to TP3 (see Figure 4). These effects were driven by larger decreases in the sleep deprived individuals than in the normal SWC group. The changes in INVF were widespread, spanning major WM tracts including the inferior fronto-occipital fasciculi, longitudinal fasciculi, thalamic radiations and the corticospinal tract. In exMD, there were widespread bilateral clusters spanning tracts as the longitudinal fasciculi, corpus callosum, thalamic radiations and the corticospinal tract. In exRD, interaction effects were observed in the inferior longitudinal fasciculi, uncinate fasciculus, anterior thalamic radiation and inferior fronto-occipital fasciculi. From TP1 to TP3, there were significant group × time interaction effects in exMD and exRD, while from TP1 to TP4 there were significant interaction effects in exMD. These effects spanned tracts such as the anterior thalamic radiation, longitudinal fasciculi, inferior fronto-occipital fasciculi and corticospinal tract. Again, these effects were driven by larger decreases in the sleep deprived individuals than in the normal SWC group. From TP2 to TP4, there were significant interaction effects in INVF, exMD and exRD in widespread WM clusters spanning major WM tracts. From TP3 to TP4, there were no significant voxel-wise group × time interaction effects. In DTI metrics, group × time interaction effects revealed a significant interaction in AD from TP2 to TP3, in MD and AD from TP1 to TP3, and in FA and RD from TP1 to TP4 (See Supplementary Material 1). Apart from the interaction in FA from TP1 to TP4, which was driven by a larger increase in the sleep deprived group, these effects were driven by a larger decrease in the sleep deprived group compared to the normal SWC group. In DKI metrics, group × time interaction effects revealed a significant interaction in MK from TP2 to TP4 in a small cluster in the right forceps minor (See Supplementary Material 2). Otherwise, there were no significant voxel-wise tests of group × time interaction effects.

**Figure 4.**
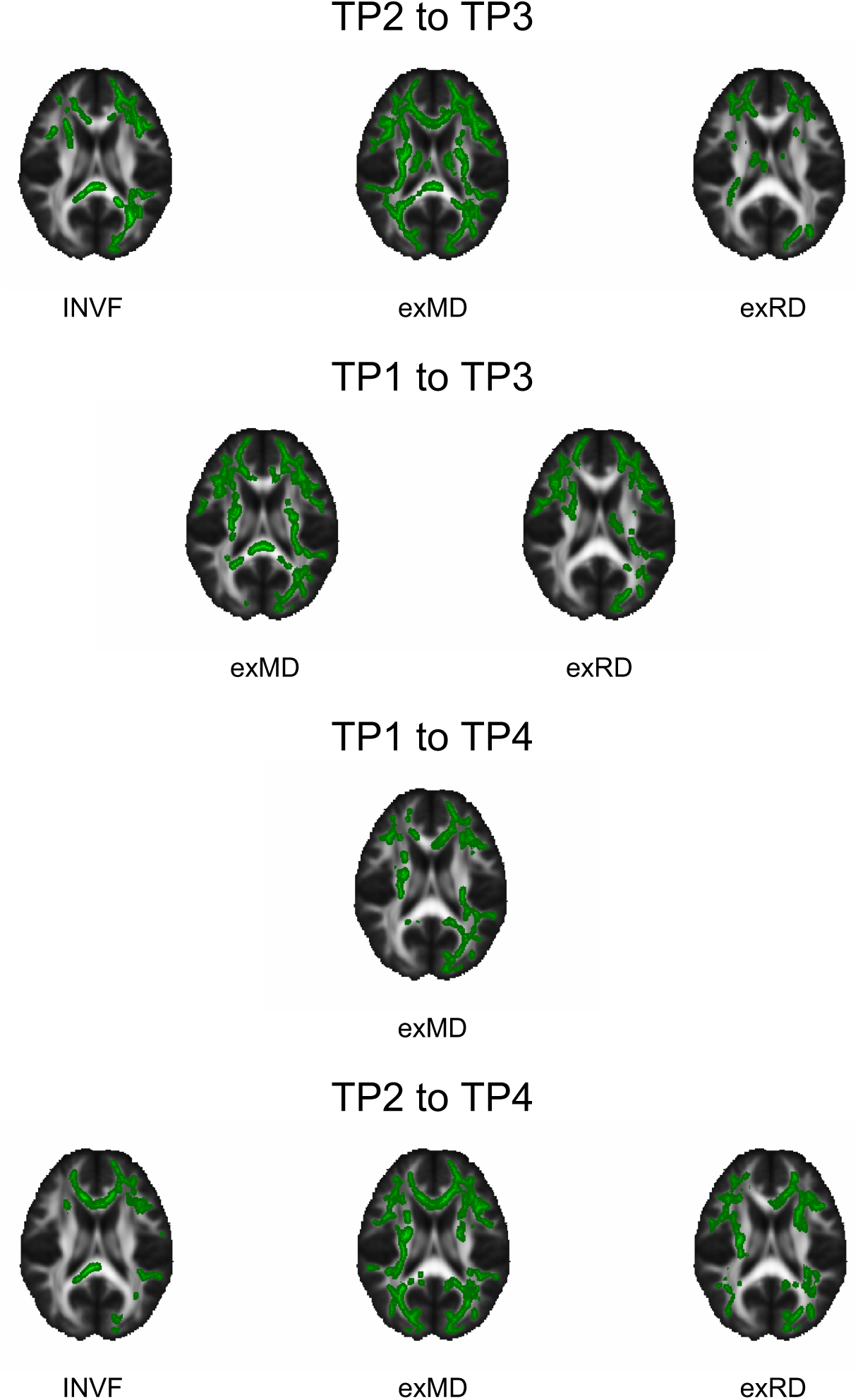
Significant voxel-wise group × time interaction effects in SMT metrics. TP; Time point. INVF; intra-neurite volume fraction. exMD; extra-neurite mean diffusivity. exRD; extra-neurite radial diffusivity. Only significant voxels are shown, corrected for family-wise error (FWE) across space and thresholded at p < 0.05.

### 3.3 Associations between diffusivity changes and sleep-wake characteristics

In the sleep deprived group from TP2 to TP4, there was a significant negative correlation between change in sleepiness and change in INVF (r = -.57, *p* = .009; see SI 3 for plots), as well as a significant positive correlation between change in sleepiness and change in exMD (r = .45, *p* = .05; see SI 3 for plots) and between change in sleepiness and change in exRD (r = .62, *p* = .004; see SI 3 for plots). However, none of these remained significant after controlling for multiple comparisons. For the remaining comparisons, there were no significant associations between change in mean diffusion values and change in sleepiness, change in alertness or any sleep-wake characteristics.

### 3.4 Changes in sleepiness and alertness

Sleepiness, as measured by KSS, and alertness, as measured by minor lapses on the PC-PVT, are presented in Figure 5. There were significant group × time interaction effects on sleepiness from TP1 to TP3 (β = 3.06, SE = .69, *p* < .001) and TP1 to TP4 (β = 2.23, SE = .72, *p* < .01), with increases in the sleep deprived group at TP3 and decreases at TP4. There were also group × time interaction effects in alertness from TP2 to TP3 (β = .366, SE = 1.36, *p* < .01), with increases of minor lapses in the sleep deprived group at TP3.

**Figure 5.**
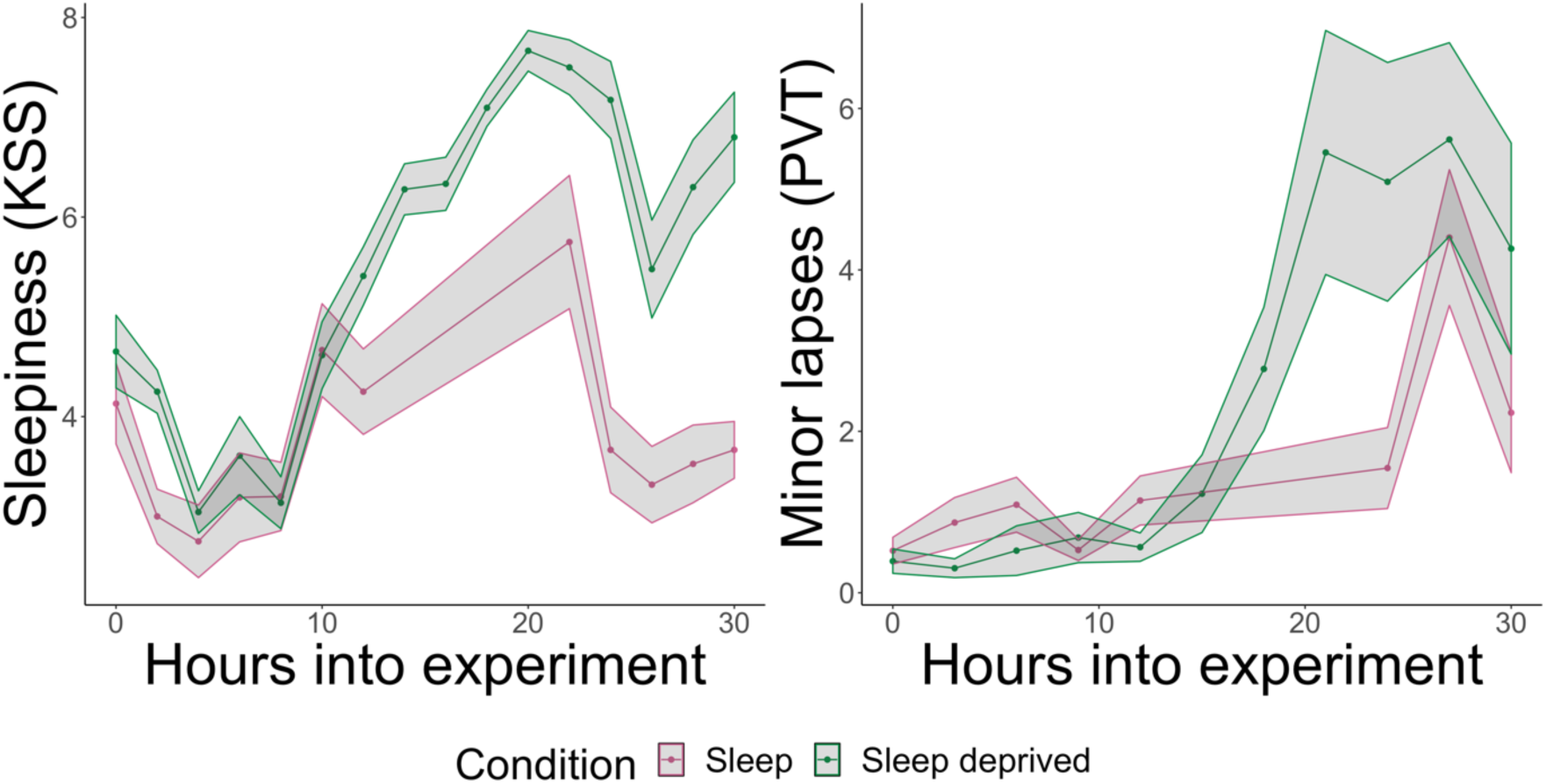
Mean (SE) of sleepiness (left) and alertness (right) across the 32 hours of sleep, wake and sleep deprivation.

## 4 Discussion

We investigated the effect of 32 hours of sleep deprivation on the human brain WM compared with a normal SWC utilising advanced diffusion modelling. We found significant group × time interaction effects in exMD, indicating that 24 and 32 hours of sleep deprivation are associated with mean diffusivity reductions in extra axonal WM. The voxel-wise tests showed that these effects span large bilateral regions of the WM skeleton, indicating global alterations across tracts. In addition, because the voxel-wise tests are more sensitive to regional changes than the global means, these tests revealed additional interaction effects in INVF and exRD. Taken together, the findings of the present study corroborate results from previous studies indicating that sleep deprivation is associated with widespread changes in WM, which differ from changes observed in a normal SWC. The use of advanced diffusion modelling allowed for estimations of microstructural substrates underlying the observed changes. The results suggest TOD effects in MRI-based measures of both intra- and extra-axonal WM microstructure, however the extra-axonal WM appeared to be more sensitive to sleep deprivation.

### 4.1 Sleep deprivation up to 24 hours

In line with our hypotheses, we found reductions in diffusivity after one night of sleep deprivation, effects that differed from those observed after a normal SWC. The interaction effect in exMD suggested a steep reduction in exMD after sleep deprivation, which was not seen after normal sleep-wake. In addition, we found interaction effects in INVF and exRD in the voxel-wise tests, with larger decreases in the sleep deprived group. These results support the view that sleep deprivation lead to alterations in WM microstructure beyond those associated with a normal SWC and TOD variations.

Our finding of interaction effects in INVF after a night of sleep deprivation compared to normal SWC are in line with our previous report of reduced FA in an independent sample of sleep deprived individuals (Elvsåshagen et al., 2015). Although INVF and FA are differentially derived metrics, they are both volume fractions sensitive to diffusion running parallel to axonal fibres (Jelescu & Budde, 2017). Furthermore, reductions in exMD and exRD in the current study might be in line with observed reductions in MD, AD and RD in the previous study that employed another MRI scanner (Elvsåshagen et al., 2015). MD, AD and RD from the DTI model are statistically derived diffusion metrics that afford indirect, less specific, measurements of brain microstructure (Jelescu & Budde, 2017), while exMD and exRD in the SMT model are more biologically specific, modelled to estimate diffusivity in the extra-axonal space (Kaden et al., 2016). The current results might thus suggest that the reductions in DTI metrics seen in the previous study are related to processes occurring in the extra-axonal space, such as cell swelling in response to neuronal activity, a possibility discussed in more detail below (Sykova & Nicholson, 2008). Importantly, we also found MD and AD interaction effects suggesting larger reductions after sleep deprivation in analyses of DTI metrics consistent with the previous study (see Supplementary Material 1 for results and figures). Together, these studies, which included different samples and MRI scanners, strongly suggest that one night of sleep deprivation is associated with changes in WM diffusion. However, other studies found no significant change in MD after sleep deprivation (Bernardi et al., 2016) or between rested wakefulness and wakefulness after 24 hours of sleep deprivation in the voxel based morphometry-derived WM volume (Demiral et al., 2019). These contrasting findings might result from substantial differences in study design, as one study combined sleep deprivation with task practice (Bernardi et al., 2016). Differences in the methods applied for measurement and estimations of WM changes may also affect results, as we relied on TBSS and mean skeleton values, while other studies have relied on regions of interest and voxel-based morphometry (Bernardi et al., 2016; Demiral et al., 2019).

We observed that changes in diffusivity after sleep deprivation were widespread, indicating that most major WM tracts might be susceptible to sleep loss. This was expected as sleep deprivation impacts a wide range of brain functions (Krause et al., 2017). The effect in tracts connecting the thalamus and brain stem was especially interesting, as brainstem nuclei play a critical role in the successful implementation of sleep state transitions (Saper & Fuller, 2017), while the anterior part of the thalamus, which project to the cingulate gyrus, hippocampus and frontal lobes, is considered to be involved in regulation of alertness (Saper, Fuller, Pedersen, Lu, & Scammell, 2010), learning and memory (Aggleton & Brown, 1999). Although we did not detect any significant associations between mean diffusivity changes and changes in alertness in this study, diffusivity changes in the relevant tracts may play a role in how sleep deprivation affects attention, learning and memory, and should be explored by future studies.

### 4.2 Sleep deprivation beyond 24 hours

The second major finding of this study was changes related to sleep deprivation beyond 24 hours. Here we found widespread interaction effects across groups after 32 hours, with opposing effects of sleep deprivation and normal SWC on mean diffusivity in extra-axonal WM. The sleep deprived group showed reductions in exMD after 32 hours, while the normal SWC group showed a small increase. In addition, the sleep deprived group showed a steep reduction in exMD from TP2 to TP3, followed by an increase from TP3 to TP4, an effect which was also seen in the normal SWC group.

If the changes in diffusion metrics observed from TP3 to TP4 were related to diurnal or circadian processes, we would expect to observe similar changes from morning to evening on the first day (TP1 to TP2). We did see similar effects in INVF, but not in exMD and exRD (see also Voldsbekk et al., 2020). One possible explanation may have to do with the diffusion metrics being sensitive and differentially affected by different aspects of sleep regulating processes. For example, changes in exMD may mainly reflect neurobiological processes that are more susceptible to sleep loss, whereas INVF could reflect changes that might be more sensitive to circadian processes. Relatedly, while diffusion changes from morning to evening may reflect neurobiological changes in WM associated with increasing sleep pressure, circadian regulation, or a combination of the two, diffusion changes after sleep deprivation are more likely to reflect mechanisms that are dominated by changes related to increasing sleep pressure. The opposing effect in exMD from TP3 to TP4, compared to TP2 to TP3, may thus reflect circadian regulation on top of increased sleep pressure, as the duration of sleep deprivation had bypassed a full cycle of the 24-hour circadian cycle.

In our analysis of associations between diffusion metric changes and sleep-wake characteristics, there were significant associations between change in subjective sleepiness and observed changes in INVF, exMD and exRD from TP2 to TP4 in the sleep deprived group. This finding supports the possibility that the observed changes in diffusion-derived brain measures after sleep deprivation might reflect processes related to increased sleep pressure. However, this finding was explorative and did not survive correction for multiple comparisons. Thus, further studies are needed to confirm this finding. Interestingly, previous studies also indicate that changes in WM microstructure after a night of sleep deprivation might be related to measures of sleepiness (Elvsåshagen et al., 2015) and visuo-motor performance (Rocklage, Williams, Pacheco, & Schnyer, 2009). Together, the current and previous associations between sleepiness and alertness and changes in several diffusion metrics encourages further exploration of whether alterations in WM microstructure may represent a neural correlate of sleepiness and alertness.

### 4.3 Potential mechanisms underlying diffusivity changes after sleep deprivation

Evidence from animal models point to several potential neurobiological mechanisms underlying the WM changes that are observed during the SWC. Transcription of genes related to macromolecule homeostasis such as protein synthesis, lipid transport (Mackiewicz et al., 2007) and lipid metabolism (Cirelli, LaVaute, & Tononi, 2005) exhibits diurnal fluctuation, and genes involved in oligodendrocyte proliferation, phospholipid synthesis and myelination are preferentially transcribed during sleep (Bellesi et al., 2013). Moreover, a recent study suggests that sleep loss may impair myelin maintenance (Bellesi et al., 2018). These findings indicate an intimate relationship between sleep regulation and maintenance of cell membranes and myelin in the brain.

While caution must be exerted in interpreting specific underlying biology on the basis of DWI (Novikov, Kiselev, & Jespersen, 2018), the SMT model hold promise to reflect measures relatable to biology. The model is validated against axonal histology (Kaden et al., 2016; Lakhani, Schilling, Xu, & Bagnato, 2020) and INVF is found to be sensitive to conditions affecting myelin and axonal integrity. Moreover, the model accommodates crossing fibres and possible orientation dispersion artefacts. A recent study also successfully applied the SMT model to estimate axon diameter index and relative volume fractions in healthy human volunteers (Fan et al., 2020). The reductions in INVF observed in the current study after sleep deprivation may thus reflect changes in such as reduced axonal integrity and breakdown of myelination maintenance, which would be consistent with findings in mice after sleep deprivation (Bellesi et al., 2018, 2013).

Based on recent years research, sleep is believed to play an important role in the distribution of soluble molecules and clearance of metabolic waste products in the central nervous system (Xie et al., 2013). This waste clearance system has been named the “glymphatic” system (Iliff et al., 2012; Taoka, Jost, Frenzel, Naganawa, & Pietsch, 2018; Wolf et al., 2019) and involves an exchange of cerebrospinal fluid (CSF) with interstitial fluid (ISF) through a system of perivascular channels formed by astroglial cells. In research animals, sleep is associated with an increase of more than 60% in interstitial space compared to wake (Xie et al., 2013), implicating that waste clearance might be more efficient during sleep due to greater availability of space for the interchange between CSF and ISF. Furthermore, preliminary MRI-derived markers of glymphatic regulation indicate larger total CSF volume during sleep compared to wake also in humans (Demiral et al., 2019). Sleep deprivation could thus cause disruption of glymphatic waste clearance of the WM parenchyma due to reduced CSF-ISF movement, which could potentially lead to accumulation of metabolites and swelling of brain structures. Such changes might contribute to the observed reductions in exMD and exRD in the sleep deprived group in our study. Moreover, the observed reductions in extra-axonal diffusivity measures might also reflect reduced ECS volume following wake-related neuronal activity, causing swelling of astrocytes, other glial cells or neurons (Sykova & Nicholson, 2008). Nevertheless, these potential mechanisms underlying changes in diffusivity after sleep deprivation remain speculative, and more research is needed to clarify the precise neurobiological mechanisms underlying the sleep deprivation-dependent diffusion changes found in the current study.

### 4.4 Limitations and future directions

Several limitations of the current results should be noted. First, the participants reported less sleep the night before the study as compared to their usual total sleep time. However, if we assume that sleep and wakefulness have opposite effects, we would expect reduced total sleep time to attenuate, rather than inflate, any waking-related diffusion changes observed in the current study. Second, although we controlled for important potential physiological confounds such as head motion, hydration level, caffeine intake, food intake and sleep habits, other factors that might differ between participants who slept and who were sleep deprived, such as respiration, cardiac pulsation, body temperature or blood pressure, were not measured and could potentially influence the MRI signal. Third, the sleep group slept in their own home during the experiment, and sleep was monitored by participants wearing an actigraph and keeping a sleep journal. However, for optimal control of exposure to *Zeitgeber* signals future studies should be conducted in a more controlled environment, such as a sleep laboratory, and sleep and sleep characteristics should be recorded using polysomnography. Fourth, although we utilised advanced diffusion modelling which can inform on biophysical tissue properties and aid interpretation of image findings compared to conventional DTI (Jelescu & Budde, 2017), the precise neural mechanisms underlying the current results remain to be elucidated by future studies.

### 4.5 Conclusion

In summary, we applied advanced diffusion modelling to study the effects of sleep deprivation on measures of human WM compared to a normal SWC, and document that sleep-wake and sleep deprivation differentially alter WM microstructure. This effect was observed in the extra-axonal water pool, indicating a particular susceptibility of extra-axonal processes to sleep loss. Although more research is needed to determine the functional significance of these WM changes and the exact underlying mechanisms, our results contribute to greater understanding of how sleep and sleep deprivation affects the human brain.

## Acknowledgements

This project was funded by a research grant from the Norwegian South-East Health Authorities (2018077, 2017090, 2015078), the Research Council of Norway (249795), the Ebbe Frøland foundation, and a research grant from Mrs. Throne-Holst.

