## Supplementary material for "Sleep and sleep deprivation differentially alter white matter microstructure: a mixed model design utilising advanced diffusion modelling"

#### 1 Results of DTI metrics

*Linear mixed models for each DTI metric.*

| | FA | MD | AD | RD |
|---|---|---|---|---|
| Group | -.05 (.30) | -.14 (.30) | -.32 (.30) | -.05 (.30) |
| TP2 | **.12$^{**}$ (.04)** | -.08$^{*}$ (.03) | -.03 (.05) | **-.10$^{**}$ (.03)** |
| TP3 | .03 (.04) | **-.10$^{**}$ (.03)** | -.12$^{**}$ (.05) | -.08$^{*}$ (.03) |
| TP4 | .02 (.04) | -.02 (.03) | -.004 (.05) | -.03 (.03) |
| Group x TP2 | .08 (.05) | -.001 (.05) | .06 (.06) | -.03 (.05) |
| Group x TP3 | .05 (.05) | -.08 (.05) | -.09 (.06) | -.07 (.05) |
| Group x TP4 | .09 (.05) | -.07 (.05) | -.04 (.06) | -.08 (.05) |
| Constant | -.05 (.21) | .14 (.21) | .21 (.21) | .10 (.21) |
| Observations | 184 | 184 | 184 | 184 |
| Log Likelihood | -16.19 | -10.91 | -46.84 | -2.77 |
| Akaike Inf. Crit. | 52.38 | 41.81 | 113.67 | 25.53 |
| Bayesian Inf. Crit. | 84.08 | 73.52 | 145.38 | 57.24 |

*Note.* Values indicate the estimated effect (β) and corresponding standard error (SE).TP; Time point. FA; fractional anisotropy. MD; mean diffusivity. AD; axial diffusivity. RD; radial diffusivity.
* indicates $p < .05$.
** indicates $p < .01$.
*** indicates $p < .001$.
**Bold** indicates significance after FDR-correction.

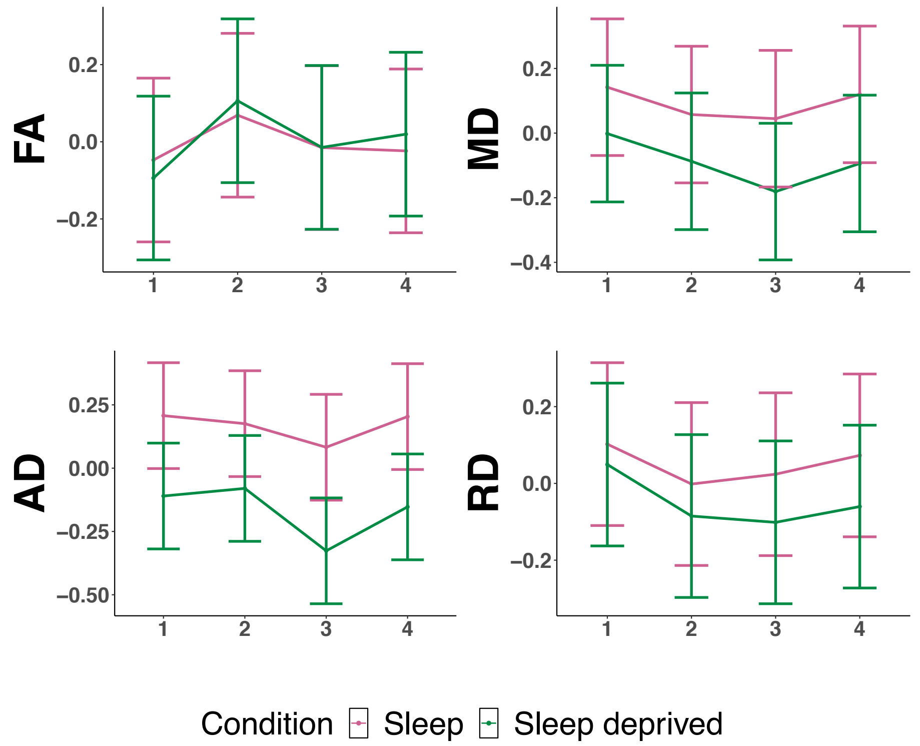


*Figure.* Estimated marginal means (y-axis) for the normal sleep-wake cycle group (pink) and sleep deprived group (green) across time points (x-axis). FA; fractional anisotropy. MD; mean diffusivity. AD; axial diffusivity. RD; radial diffusivity.

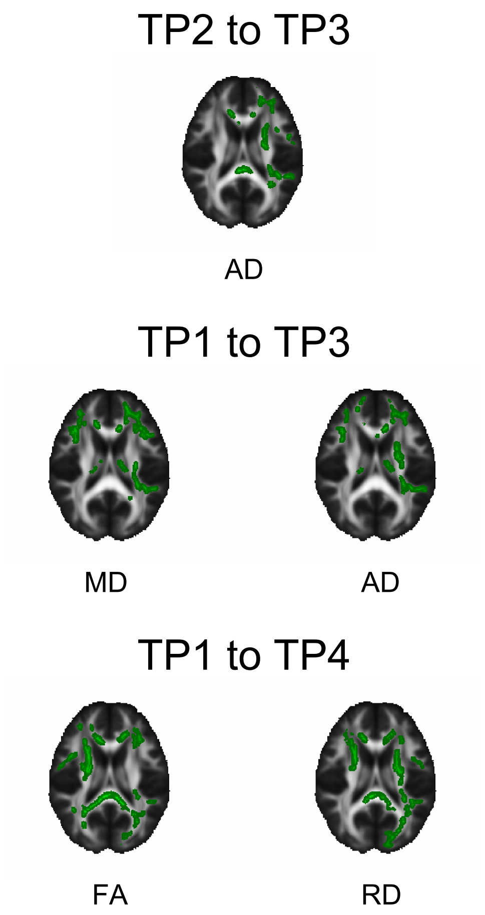


*Figure.* Significant voxel-wise group × time interaction effects in DTI metrics. TP; Time point. FA; fractional anisotropy. MD; mean diffusivity. AD; axial diffusivity. RD; radial diffusivity. Only significant voxels are shown, corrected for family-wise error (FWE) across space and thresholded at $p < 0.05$.

#### 2 Results of DKI metrics

*Linear mixed models for each diffusion metric.*

| | MK | AK | RK |
|---|---|---|---|
| Group | .03 (.30) | .16 (.30) | -.06 (.30) |
| TP2 | .09 (.05) | **.13$^{**}$ (.04)** | .06 (.07) |
| TP3 | .01 (.05) | .09$^{*}$ (.04) | -.03 (.07) |
| TP4 | .07 (.05) | .06 (.04) | .05 (.07) |
| Group x TP2 | .06 (.07) | -.04 (.05) | .13 (.10) |
| Group x TP3 | .05 (.07) | .03 (.05) | .09 (.10) |
| Group x TP4 | -.04 (.07) | .02 (.05) | -.04 (.10) |
| Constant | -.07 (.21) | -.15 (.21) | -.01 (.21) |
| Observations | 184 | 184 | 184 |
| Log Likelihood | -62.04 | -21.13 | -108.45 |
| Akaike Inf. Crit. | 144.08 | 62.27 | 236.89 |
| Bayesian Inf. Crit. | 175.78 | 93.97 | 268.60 |

*Note.* Values indicate the estimated effect (β) and corresponding standard error (SE).TP; Time point. MK; mean kurtosis. AK; axial kurtosis. RK; radial kurtosis.
* indicates $p < .05$.
** indicates $p < .01$.
*** indicates $p < .001$.
**Bold** indicates significance after FDR-correction.

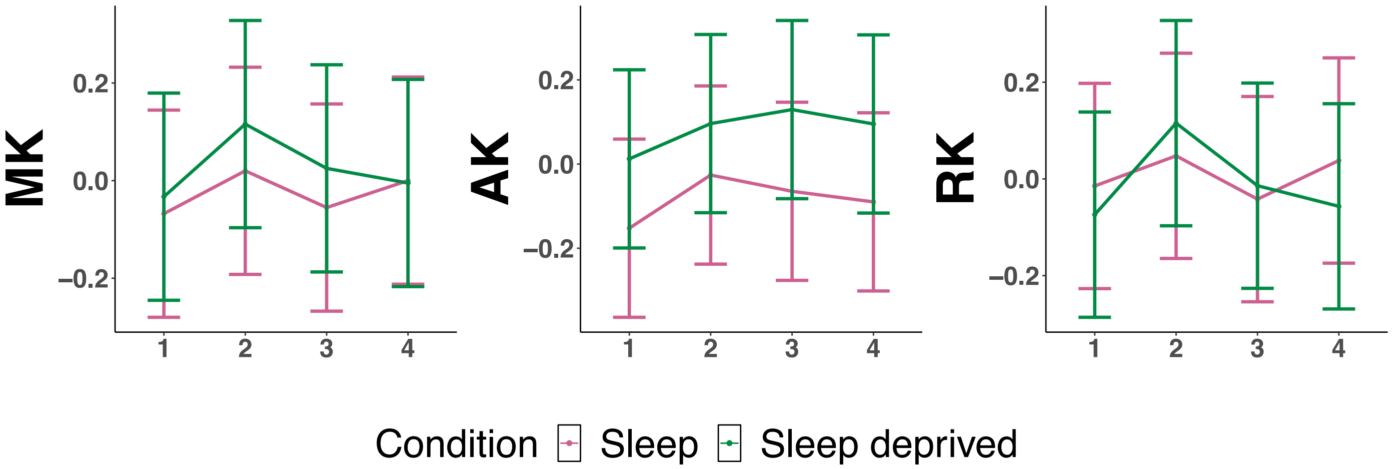


*Figure.* Estimated marginal means (y-axis) for the normal sleep-wake cycle group (pink) and sleep deprived group (green) across time points (x-axis). MK; mean kurtosis. AK; axial kurtosis. RK; radial kurtosis.

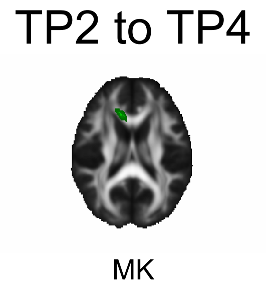


*Figure.* Significant voxel-wise group × time interaction effects in DKI metrics. TP; Time point. MK; mean kurtosis. Only significant voxels are shown, corrected for family-wise error (FWE) across space and thresholded at $p < 0.05$.

#### 3 Correlation plots of associations between group by time interaction effects in diffusion metrics and sleep-wake characteristics

##### Changes from the evening to the next morning (TP2 versus TP3)

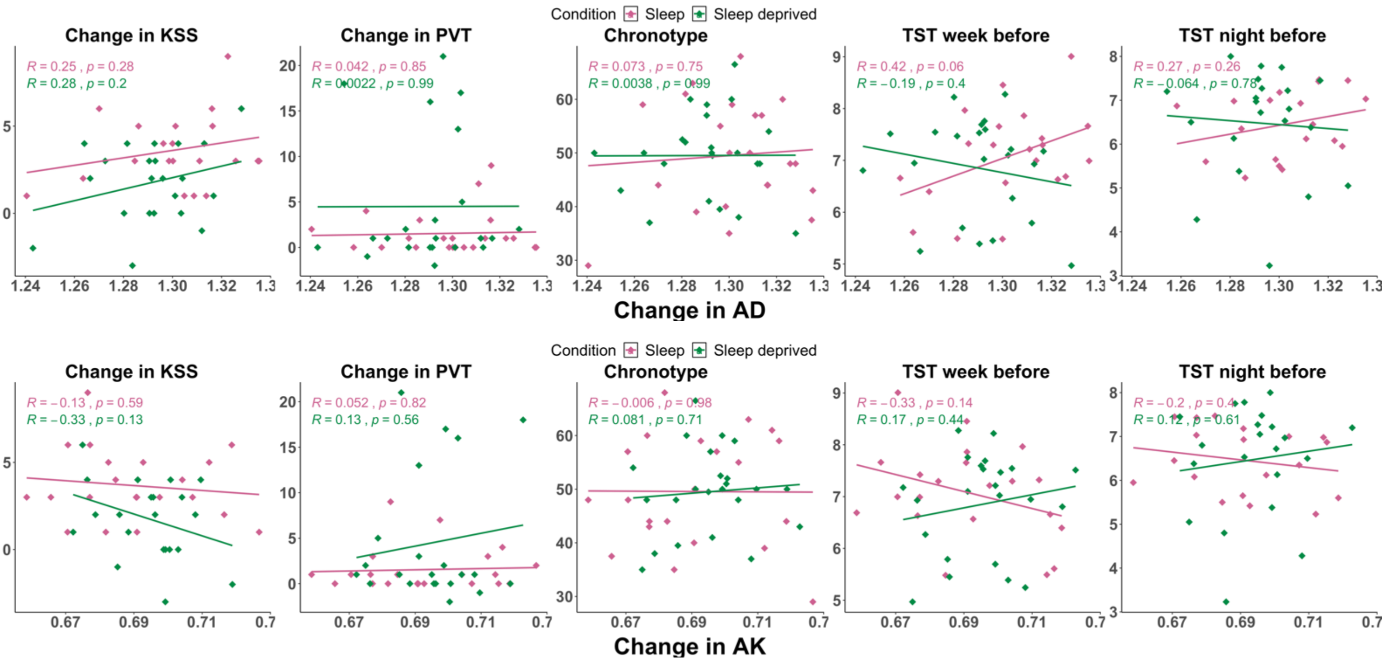

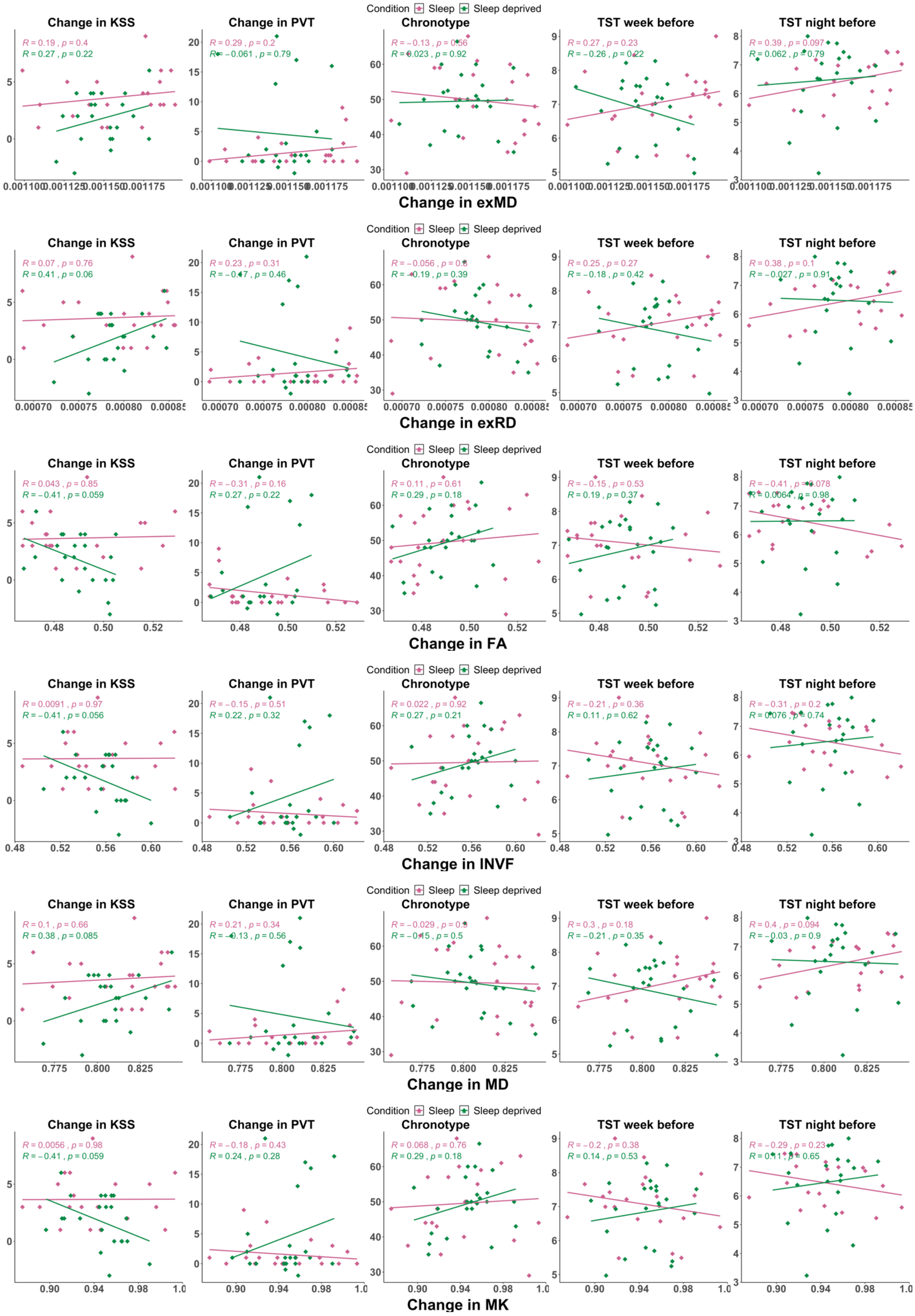
Condition Sleep Sleep deprived
Change in KSS
Change in PVT
Chronotype
TST week before
TST night before
Change in exMD
Change in exRD
Change in FA
Change in INVF
Change in MD
Change in MK

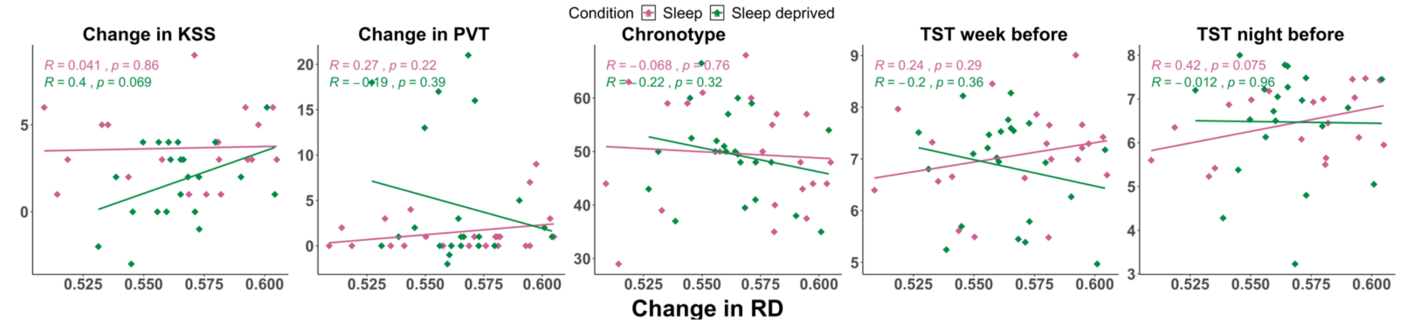


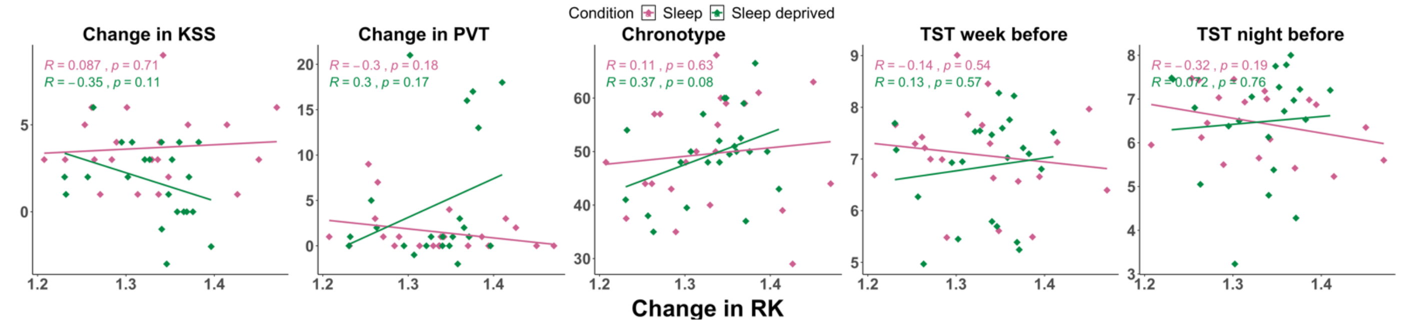


### Changes from the first to the second morning (TP1 versus TP3)

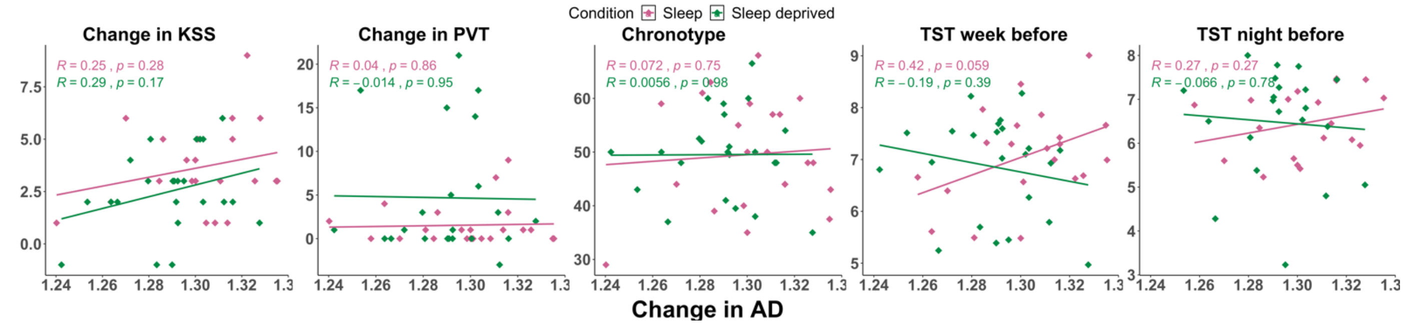


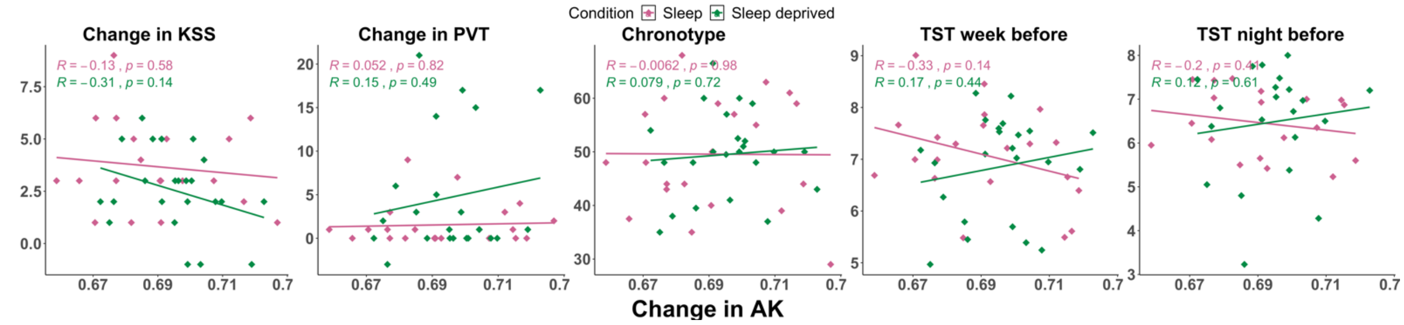


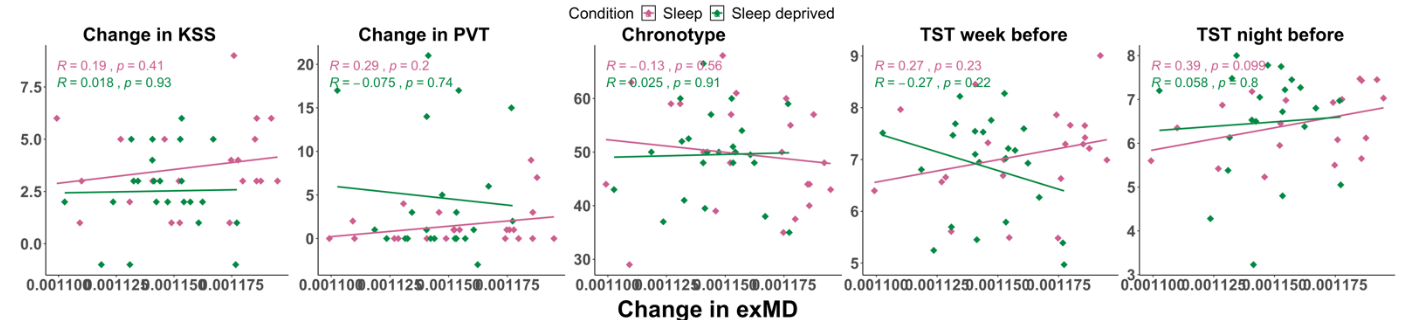

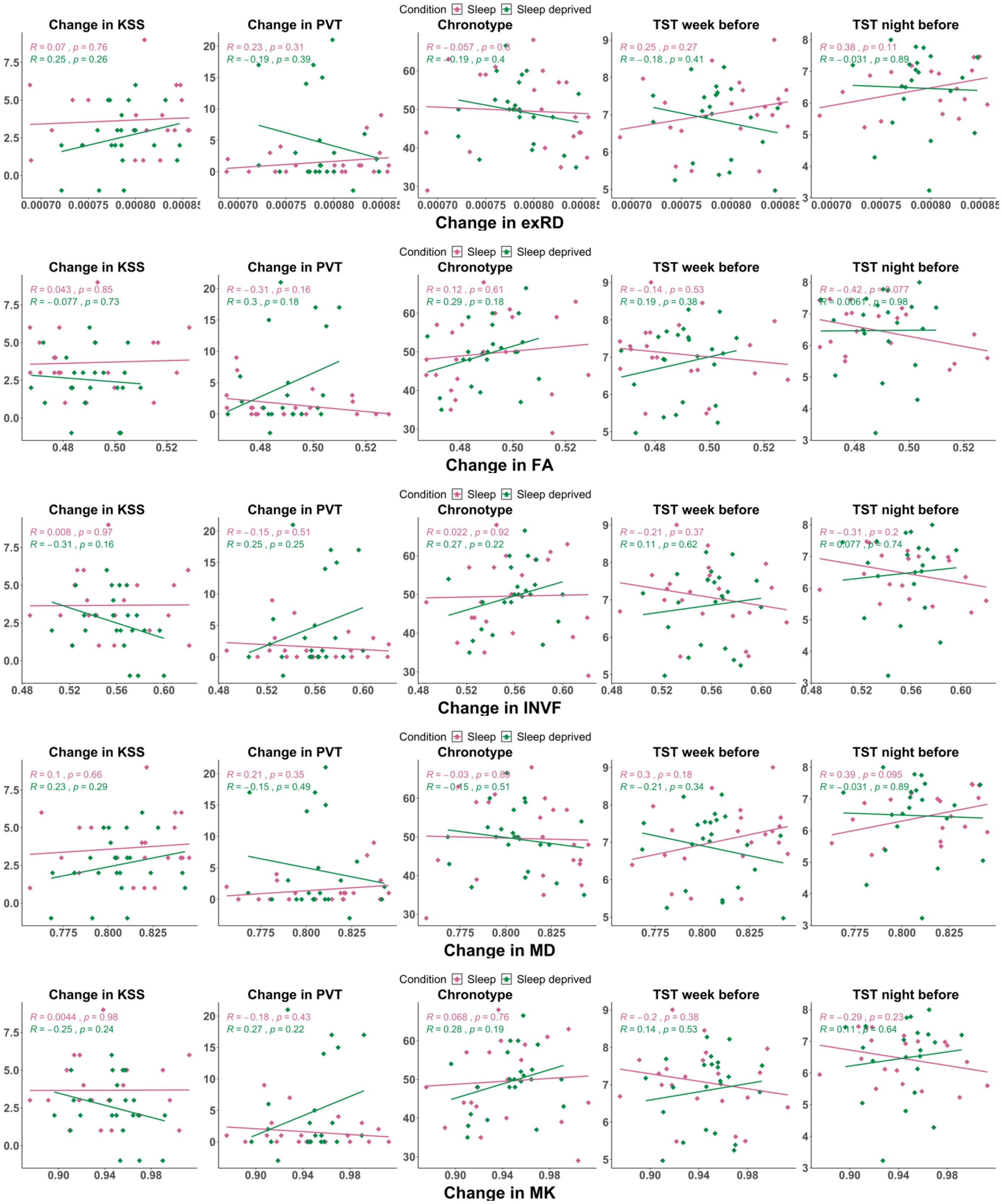
Condition Sleep Sleep deprived
Change in KSS
R = 0.07 , p = 0.76
R = 0.25 , p = 0.26
Change in PVT
R = 0.23 , p = 0.31
R = −0.19 , p = 0.39
Chronotype
R = −0.19 , p = 0.4
TST week before
R = 0.25 , p = 0.27
R = −0.18 , p = 0.41
TST night before
R = 0.38 , p = 0.11
R = −0.031 , p = 0.89
Change in exRD
Condition Sleep Sleep deprived
Change in KSS
R = 0.043 , p = 0.85
R = −0.077 , p = 0.73
Change in PVT
R = −0.31 , p = 0.16
R = 0.3 , p = 0.18
Chronotype
R = 0.12 , p = 0.61
R = 0.29 , p = 0.18
TST week before
R = −0.14 , p = 0.53
R = 0.19 , p = 0.38
TST night before
R = −0.42 , p = 0.077
R = 0.0061 , p = 0.98
Change in FA
Condition Sleep Sleep deprived
Change in KSS
R = 0.008 , p = 0.97
R = −0.31 , p = 0.16
Change in PVT
R = −0.15 , p = 0.51
R = 0.25 , p = 0.25
Chronotype
R = 0.022 , p = 0.92
R = 0.27 , p = 0.22
TST week before
R = −0.21 , p = 0.37
R = 0.11 , p = 0.62
TST night before
R = −0.31 , p = 0.2
R = 0.077 , p = 0.74
Change in INVF
Condition Sleep Sleep deprived
Change in KSS
R = 0.1 , p = 0.66
R = 0.23 , p = 0.29
Change in PVT
R = 0.21 , p = 0.35
R = −0.15 , p = 0.49
Chronotype
R = −0.15 , p = 0.51
TST week before
R = 0.3 , p = 0.18
R = −0.21 , p = 0.34
TST night before
R = 0.39 , p = 0.095
R = −0.031 , p = 0.89
Change in MD
Condition Sleep Sleep deprived
Change in KSS
R = 0.0044 , p = 0.98
R = −0.25 , p = 0.24
Change in PVT
R = −0.18 , p = 0.43
R = 0.27 , p = 0.22
Chronotype
R = 0.068 , p = 0.76
R = 0.28 , p = 0.19
TST week before
R = −0.2 , p = 0.38
R = 0.14 , p = 0.53
TST night before
R = −0.29 , p = 0.23
R = 0.11 , p = 0.64
Change in MK

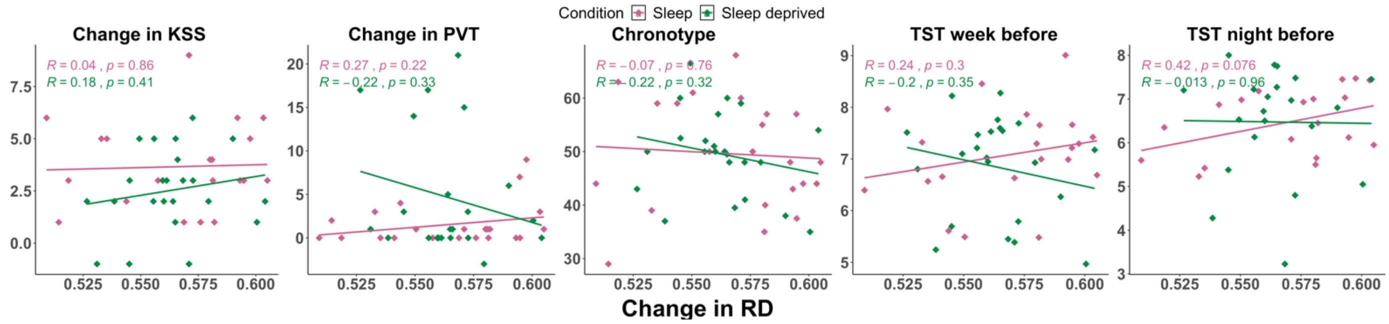


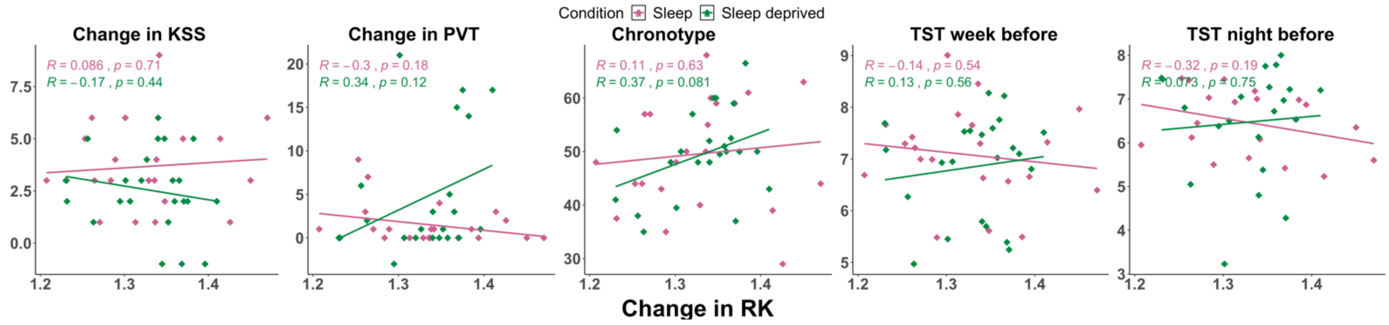


#### Changes from the first morning to the next day afternoon (TP1 versus TP4)

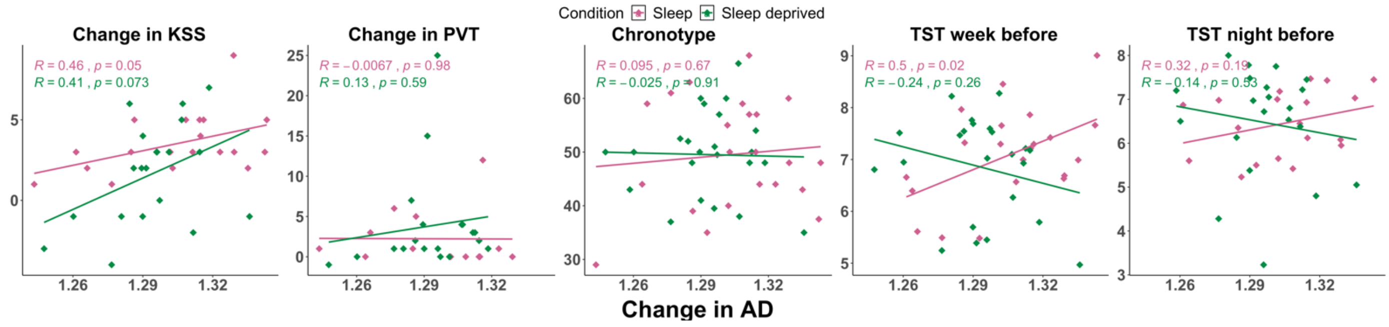


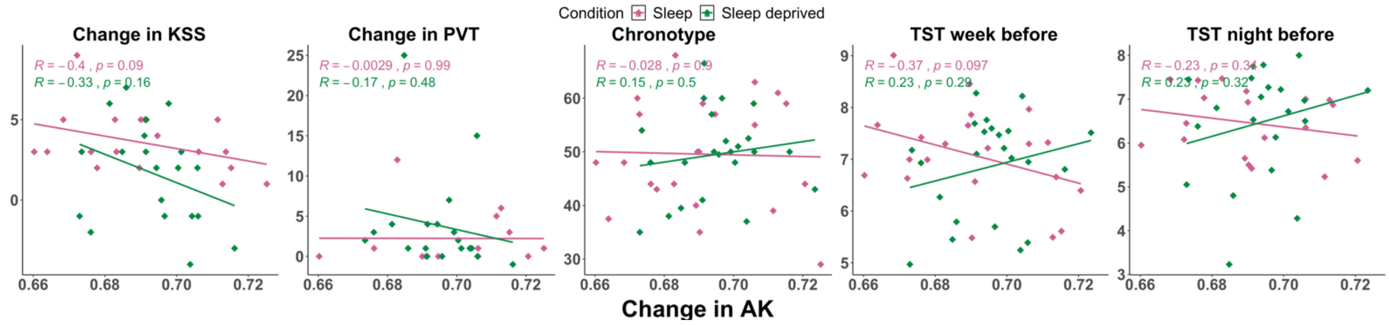

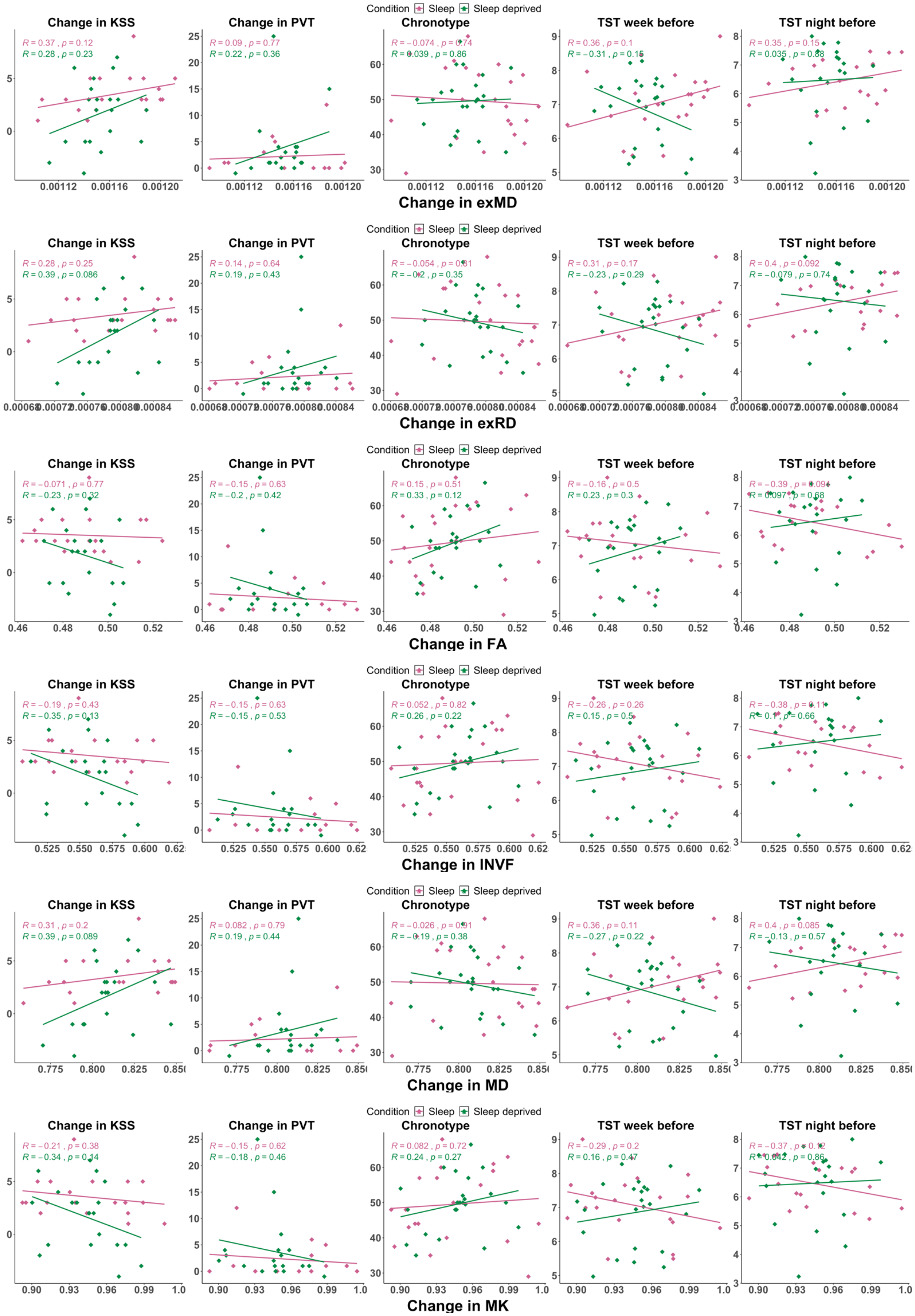
Condition Sleep Sleep deprived
Change in KSS
R = 0.37 , p = 0.12
R = 0.28 , p = 0.23
Change in PVT
R = 0.09 , p = 0.77
R = 0.22 , p = 0.36
Chronotype
R = −0.074 , p = 0.74
R = 0.039 , p = 0.86
TST week before
R = 0.36 , p = 0.1
R = −0.31 , p = 0.15
TST night before
R = 0.35 , p = 0.15
R = 0.035 , p = 0.88
Change in exMD
Condition Sleep Sleep deprived
Change in KSS
R = 0.28 , p = 0.25
R = 0.39 , p = 0.086
Change in PVT
R = 0.14 , p = 0.64
R = 0.19 , p = 0.43
Chronotype
R = −0.054 , p = 0.81
R = −0.2 , p = 0.35
TST week before
R = 0.31 , p = 0.17
R = −0.23 , p = 0.29
TST night before
R = 0.4 , p = 0.092
R = −0.079 , p = 0.74
Change in exRD
Condition Sleep Sleep deprived
Change in KSS
R = −0.071 , p = 0.77
R = −0.23 , p = 0.32
Change in PVT
R = −0.15 , p = 0.63
R = −0.2 , p = 0.42
Chronotype
R = 0.15 , p = 0.51
R = 0.33 , p = 0.12
TST week before
R = −0.16 , p = 0.5
R = 0.23 , p = 0.3
TST night before
R = −0.39 , p = 0.094
R = 0.097 , p = 0.68
Change in FA
Condition Sleep Sleep deprived
Change in KSS
R = −0.19 , p = 0.43
R = −0.35 , p = 0.13
Change in PVT
R = −0.15 , p = 0.63
R = −0.15 , p = 0.53
Chronotype
R = 0.052 , p = 0.82
R = 0.26 , p = 0.22
TST week before
R = −0.26 , p = 0.26
R = 0.15 , p = 0.5
TST night before
R = −0.38 , p = 0.11
R = 0.1 , p = 0.66
Change in INVF
Condition Sleep Sleep deprived
Change in KSS
R = 0.31 , p = 0.2
R = 0.39 , p = 0.089
Change in PVT
R = 0.082 , p = 0.79
R = 0.19 , p = 0.44
Chronotype
R = −0.026 , p = 0.91
R = −0.19 , p = 0.38
TST week before
R = 0.36 , p = 0.11
R = −0.27 , p = 0.22
TST night before
R = 0.4 , p = 0.085
R = −0.13 , p = 0.57
Change in MD
Condition Sleep Sleep deprived
Change in KSS
R = −0.21 , p = 0.38
R = −0.34 , p = 0.14
Change in PVT
R = −0.15 , p = 0.62
R = −0.18 , p = 0.46
Chronotype
R = 0.082 , p = 0.72
R = 0.24 , p = 0.27
TST week before
R = −0.29 , p = 0.2
R = 0.16 , p = 0.47
TST night before
R = −0.37 , p = 0.12
R = 0.042 , p = 0.86
Change in MK

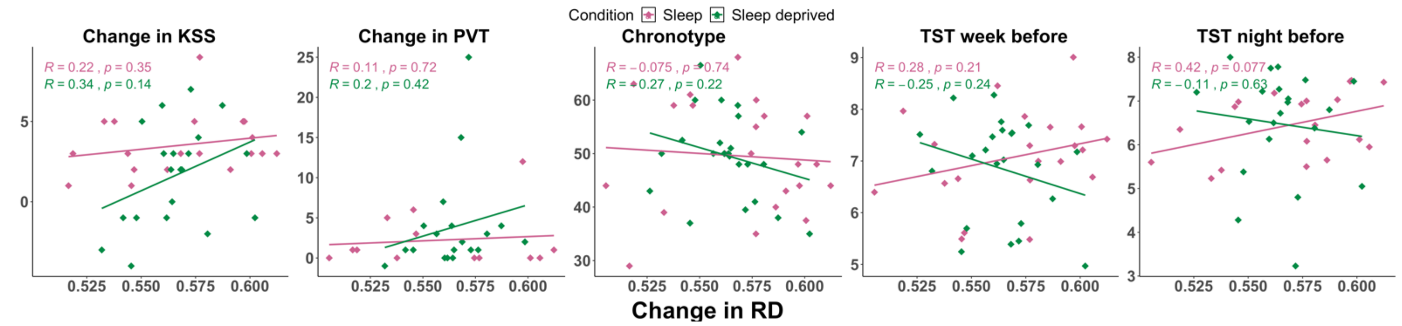


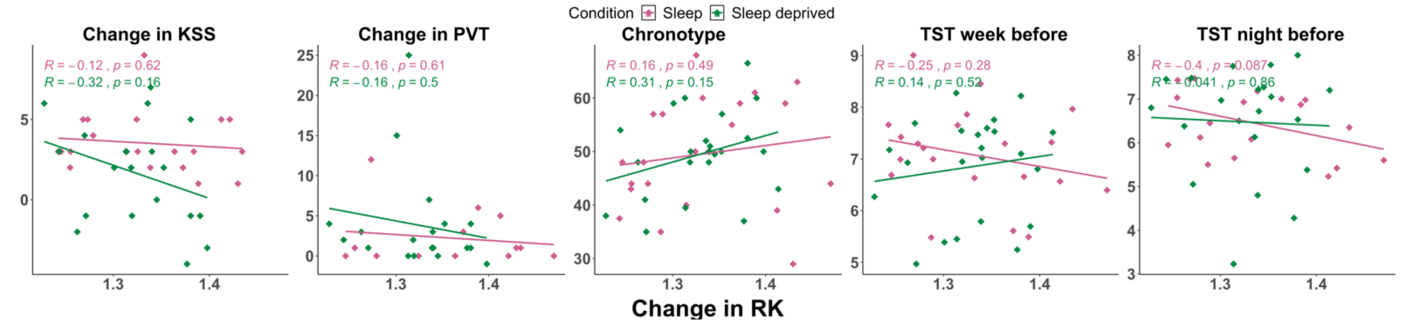


### Changes from the evening to the next day afternoon (TP2 versus TP4)

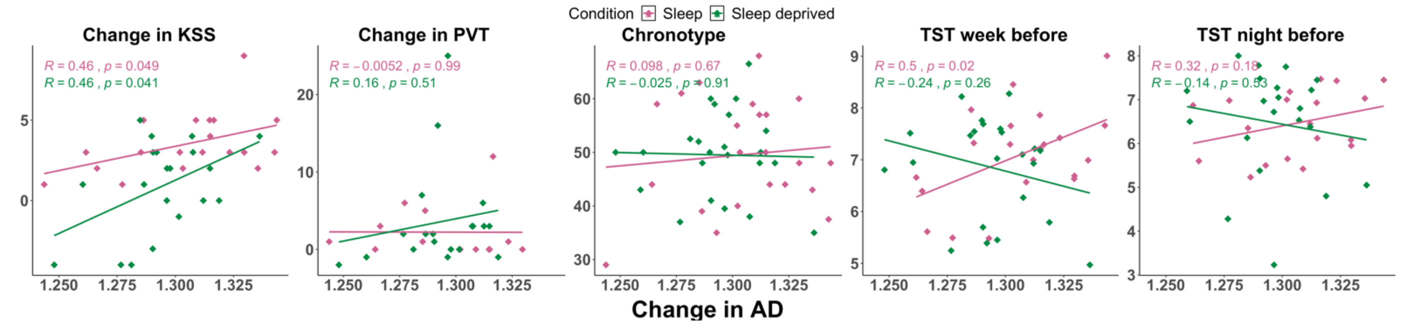


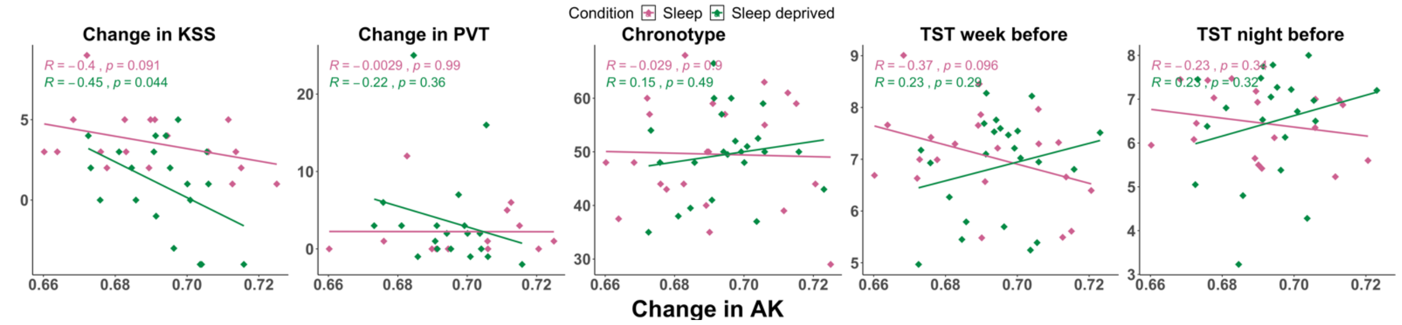


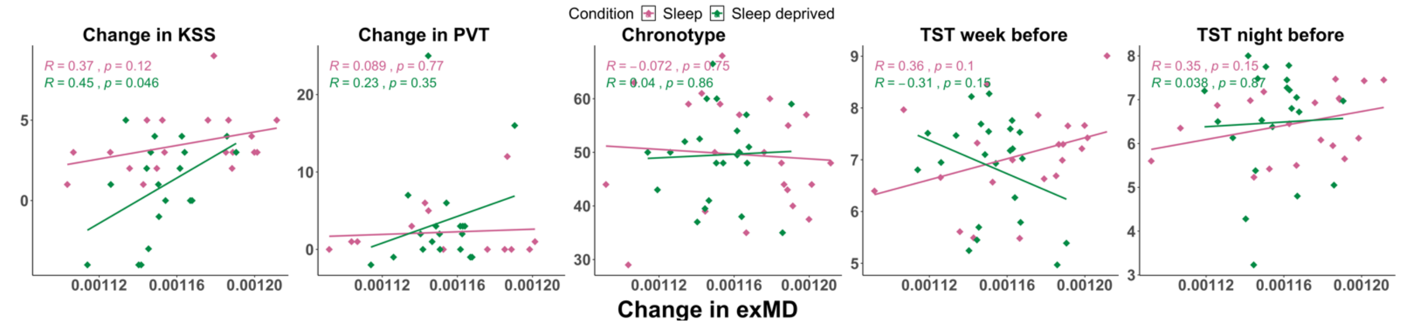

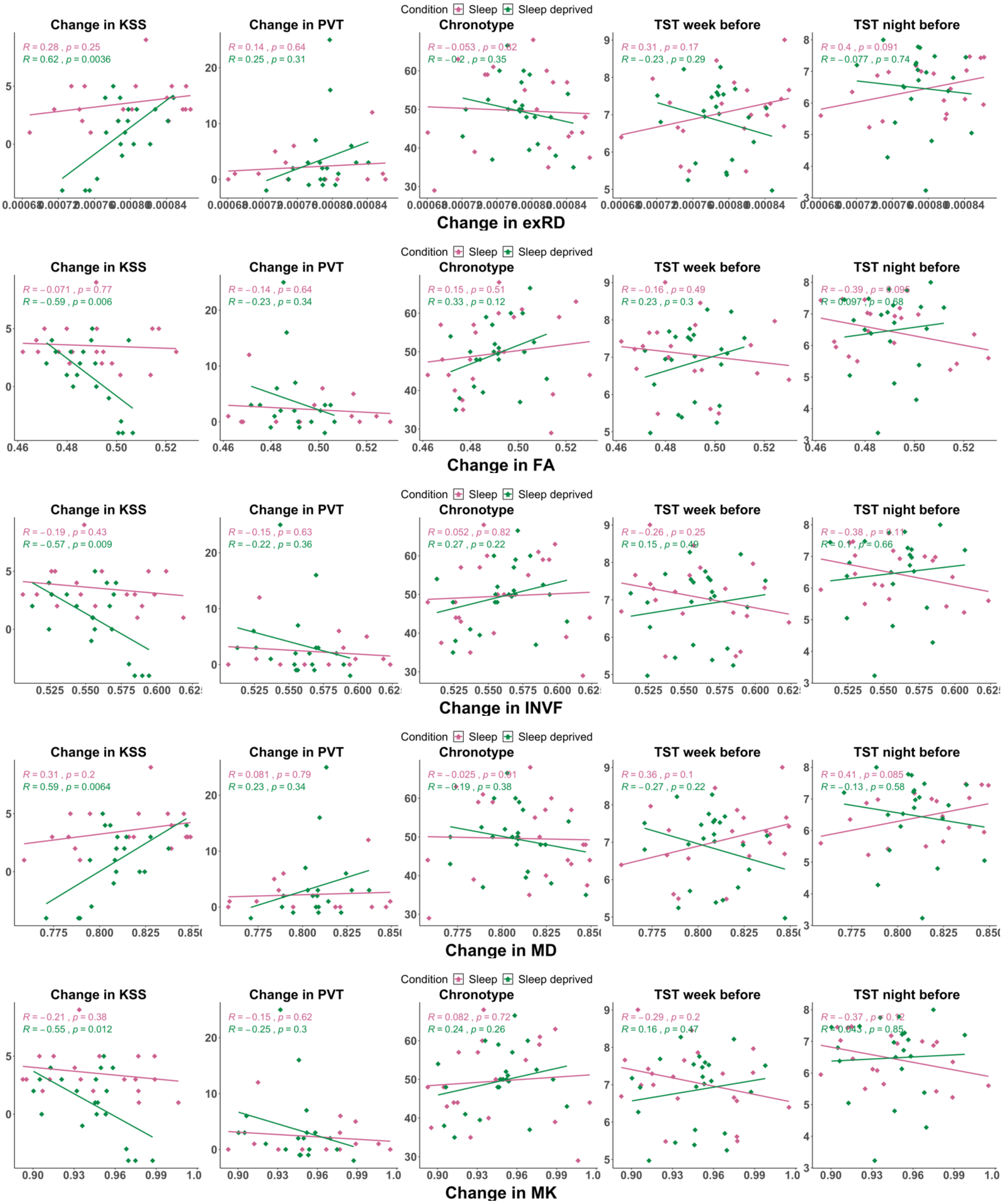

Condition Sleep Sleep deprived
Change in KSS
Change in PVT
Chronotype
TST week before
TST night before
R = 0.28 , p = 0.25
R = 0.62 , p = 0.0036
R = 0.14 , p = 0.64
R = 0.25 , p = 0.31
R = −0.053 , p = 0.82
R = 0.31 , p = 0.17
R = −0.23 , p = 0.29
R = 0.4 , p = 0.091
R = −0.077 , p = 0.74
Change in exRD
R = −0.071 , p = 0.77
R = −0.59 , p = 0.006
R = −0.14 , p = 0.64
R = −0.23 , p = 0.34
R = 0.15 , p = 0.51
R = 0.33 , p = 0.12
R = −0.16 , p = 0.49
R = 0.23 , p = 0.3
R = −0.39 , p = 0.095
R = 0.097 , p = 0.68
Change in FA
R = −0.19 , p = 0.43
R = −0.57 , p = 0.009
R = −0.15 , p = 0.63
R = −0.22 , p = 0.36
R = 0.052 , p = 0.82
R = 0.27 , p = 0.22
R = −0.26 , p = 0.25
R = 0.15 , p = 0.49
R = −0.38 , p = 0.11
R = 0.1 , p = 0.66
Change in INVF
R = 0.31 , p = 0.2
R = 0.59 , p = 0.0064
R = 0.081 , p = 0.79
R = 0.23 , p = 0.34
R = −0.025 , p = 0.91
R = −0.19 , p = 0.38
R = 0.36 , p = 0.1
R = −0.27 , p = 0.22
R = 0.41 , p = 0.085
R = −0.13 , p = 0.58
Change in MD
R = −0.21 , p = 0.38
R = −0.55 , p = 0.012
R = −0.15 , p = 0.62
R = −0.25 , p = 0.3
R = 0.082 , p = 0.72
R = 0.24 , p = 0.26
R = −0.29 , p = 0.2
R = 0.16 , p = 0.47
R = −0.37 , p = 0.12
R = 0.043 , p = 0.85
Change in MK

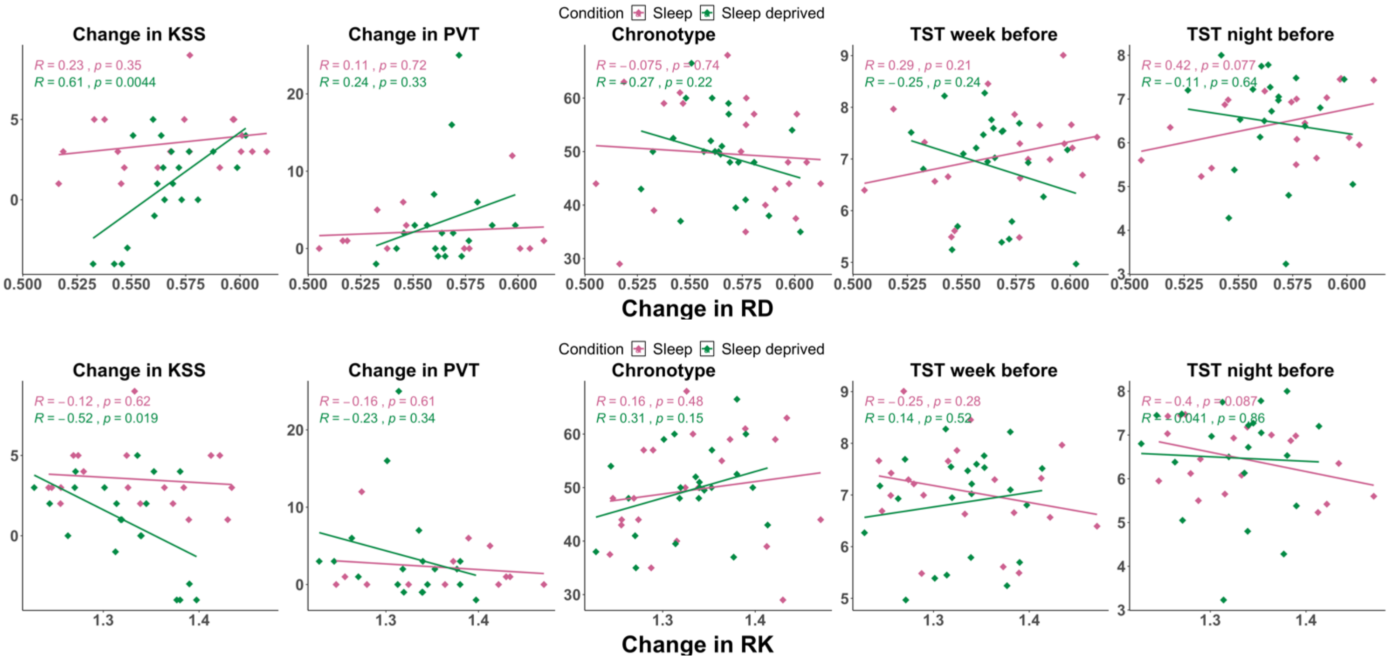


#### Changes from the second morning to the same day afternoon (TP3 versus TP4)

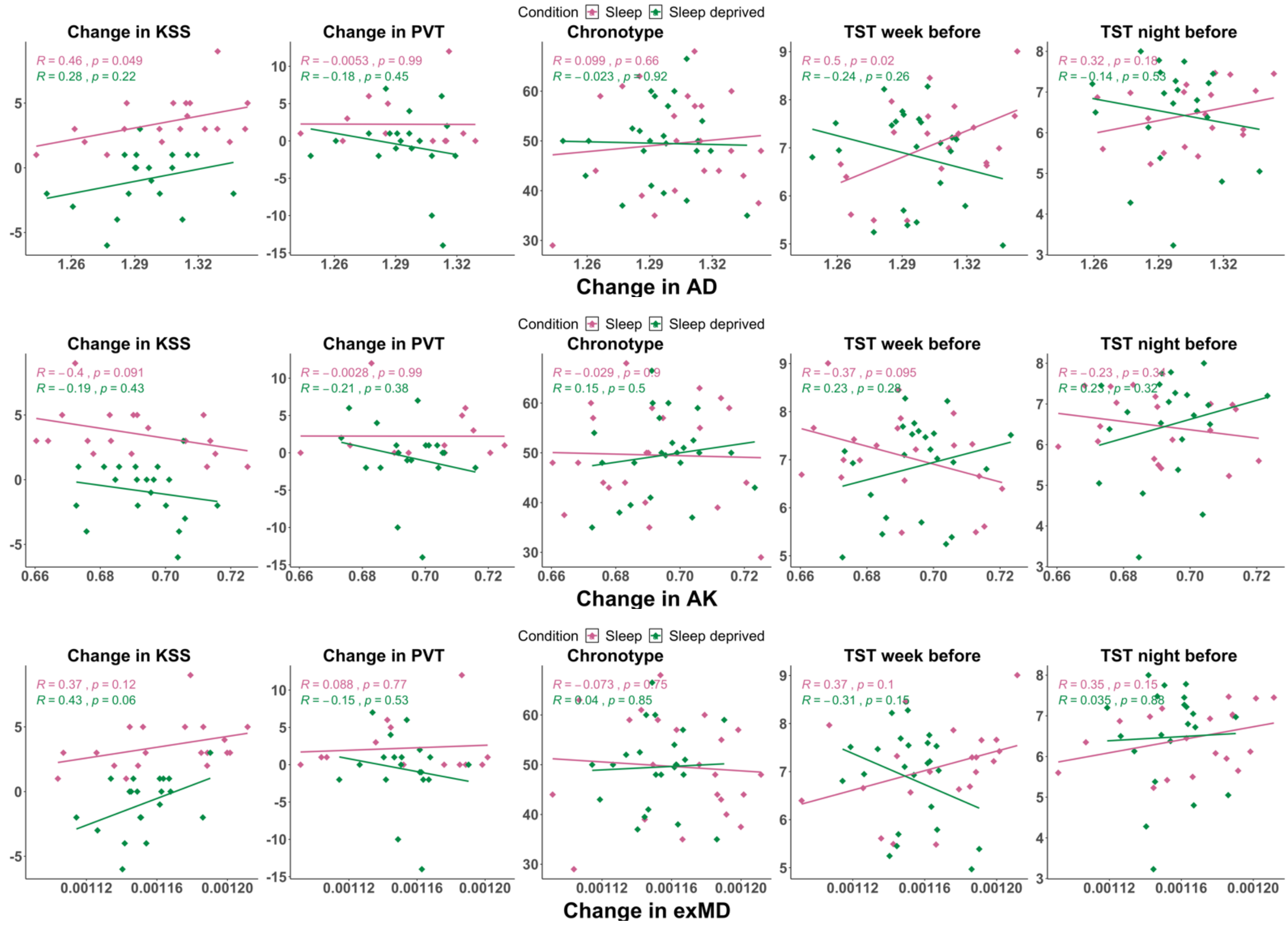

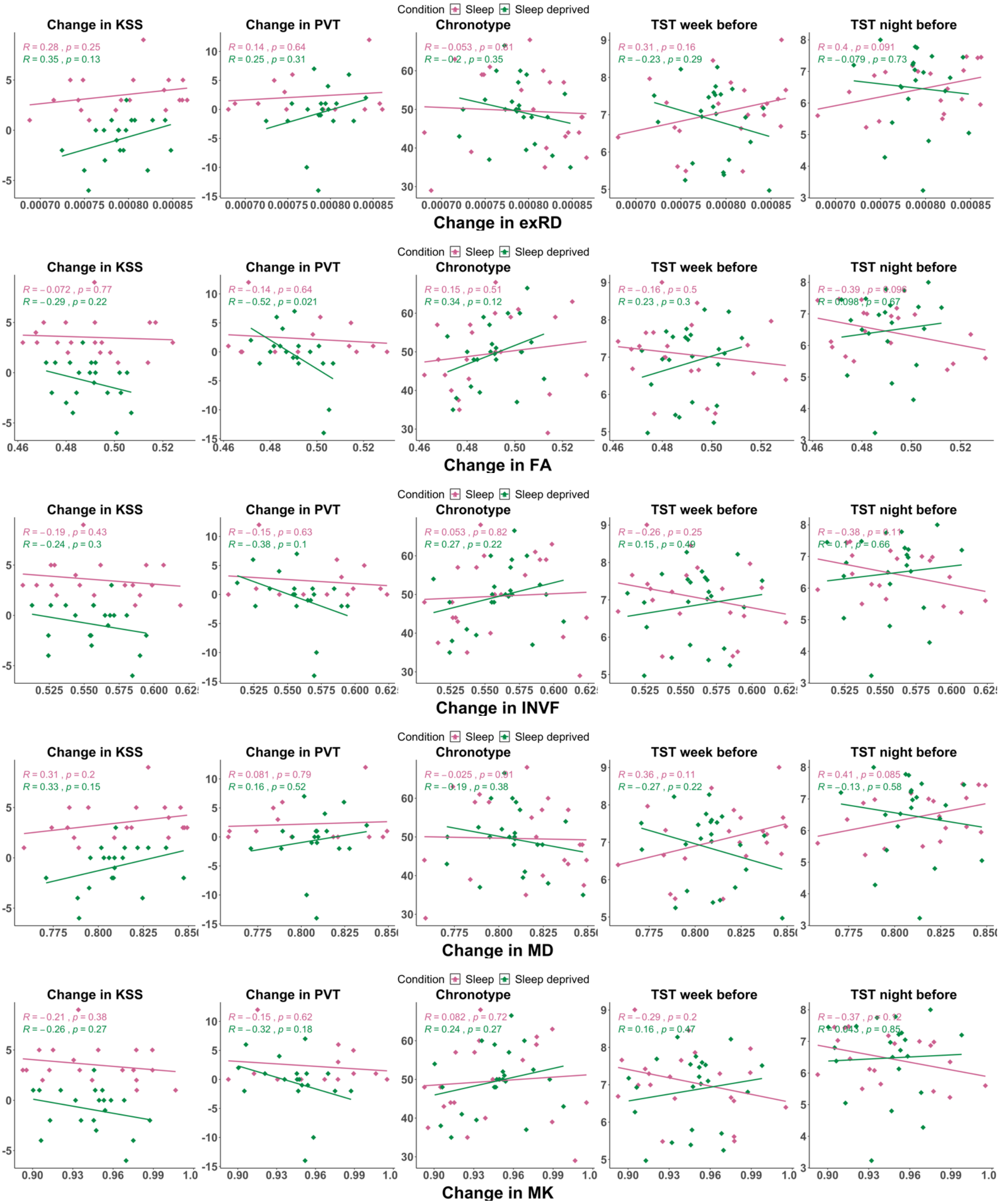

Condition Sleep Sleep deprived
Change in KSS
Change in PVT
Chronotype
TST week before
TST night before
R = 0.28 , p = 0.25
R = 0.35 , p = 0.13
R = 0.14 , p = 0.64
R = 0.25 , p = 0.31
R = −0.053 , p = 0.81
R = 0.35
R = 0.31 , p = 0.16
R = −0.23 , p = 0.29
R = 0.4 , p = 0.091
R = −0.079 , p = 0.73
Change in exRD
R = −0.072 , p = 0.77
R = −0.29 , p = 0.22
R = −0.14 , p = 0.64
R = −0.52 , p = 0.021
R = 0.15 , p = 0.51
R = 0.34 , p = 0.12
R = −0.16 , p = 0.5
R = 0.23 , p = 0.3
R = −0.39 , p = 0.096
R = 0.098 , p = 0.67
Change in FA
R = −0.19 , p = 0.43
R = −0.24 , p = 0.3
R = −0.15 , p = 0.63
R = −0.38 , p = 0.1
R = 0.053 , p = 0.82
R = 0.27 , p = 0.22
R = −0.26 , p = 0.25
R = 0.15 , p = 0.49
R = −0.38 , p = 0.11
R = 0.1 , p = 0.66
Change in INVF
R = 0.31 , p = 0.2
R = 0.33 , p = 0.15
R = 0.081 , p = 0.79
R = 0.16 , p = 0.52
R = −0.025 , p = 0.91
R = −0.19 , p = 0.38
R = 0.36 , p = 0.11
R = −0.27 , p = 0.22
R = 0.41 , p = 0.085
R = −0.13 , p = 0.58
Change in MD
R = −0.21 , p = 0.38
R = −0.26 , p = 0.27
R = −0.15 , p = 0.62
R = −0.32 , p = 0.18
R = 0.082 , p = 0.72
R = 0.24 , p = 0.27
R = −0.29 , p = 0.2
R = 0.16 , p = 0.47
R = −0.37
R = 0.043 , p = 0.85
Change in MK

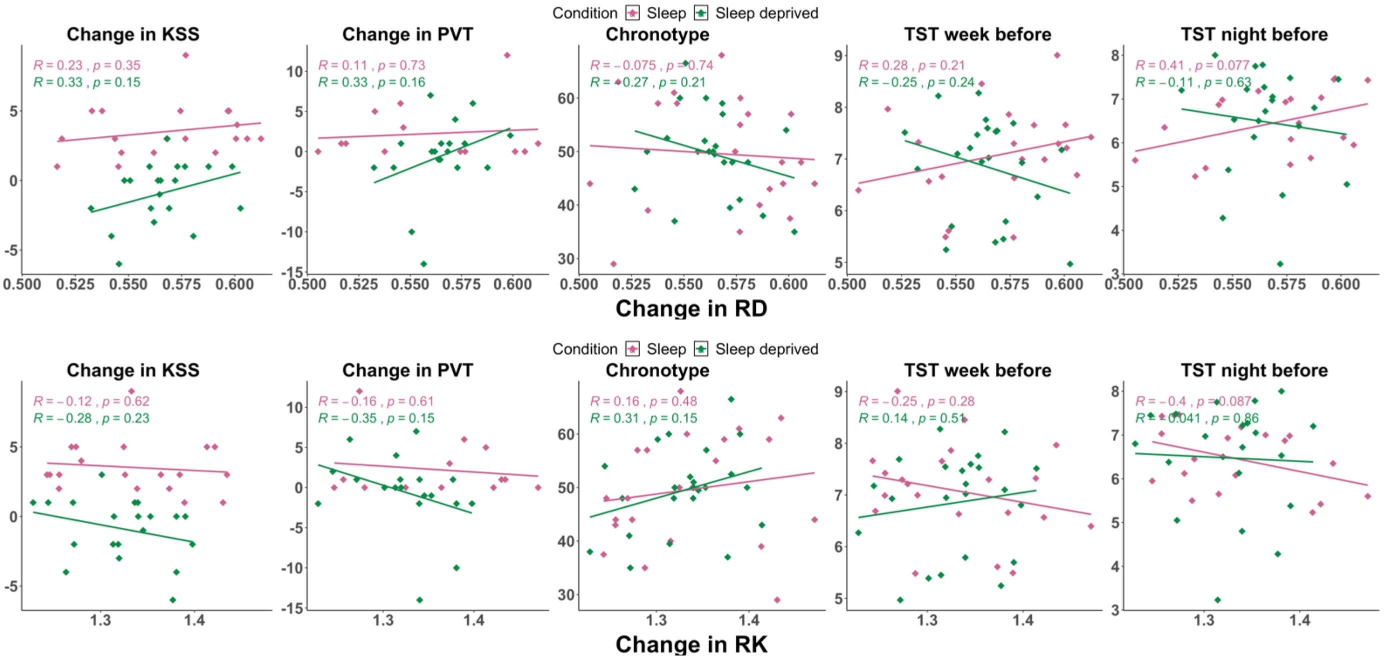
Condition Sleep Sleep deprived
Change in KSS
R = 0.23 , p = 0.35
R = 0.33 , p = 0.15
Change in PVT
R = 0.11 , p = 0.73
R = 0.33 , p = 0.16
Chronotype
R = −0.075 , p = 0.74
R = −0.27 , p = 0.21
TST week before
R = 0.28 , p = 0.21
R = −0.25 , p = 0.24
TST night before
R = 0.41 , p = 0.077
R = −0.11 , p = 0.63
Change in RD
Condition Sleep Sleep deprived
Change in KSS
R = −0.12 , p = 0.62
R = −0.28 , p = 0.23
Change in PVT
R = −0.16 , p = 0.61
R = −0.35 , p = 0.15
Chronotype
R = 0.16 , p = 0.48
R = 0.31 , p = 0.15
TST week before
R = −0.25 , p = 0.28
R = 0.14 , p = 0.51
TST night before
R = −0.4 , p = 0.087
R = 0.041 , p = 0.86
Change in RK
